# Sparse learning for scalable phylogenetic network inference

**DOI:** 10.1101/2025.11.16.688704

**Authors:** Ulises Rosas-Puchuri, Nathan Kolbow, Claudia Solís-Lemus, Sina Khanmohammadi, Betancur-R. Ricardo

**Affiliations:** School of Biological Sciences, The University of Oklahoma, Norman, Oklahoma, USA; Wisconsin Institute for Discovery, University of Wisconsin-Madison, Madison, Wisconsin, USA; Department of Statistics, University of Wisconsin-Madison, Madison, Wisconsin, USA; Department of Plant Pathology, University of Wisconsin-Madison, Madison, Wisconsin, USA; School of Computer Science, The University of Oklahoma, Norman, Oklahoma, USA; Data Science and Analytics Institute, The University of Oklahoma, Norman, Oklahoma, USA; Scripps Institution of Oceanography, University of California San Diego, La Jolla, California, USA

**Keywords:** Supervised machine learning, LASSO regression, Level-1 phylogenetic networks, Dimensionality reduction, Regression trees

## Abstract

Phylogenetic networks account for signals of hybridization, reticulation, and gene flow, and provide an opportunity to analyse species evolution from a more complex perspective than bifurcating phylogenetic trees. However, even the fastest algorithms for inferring these networks face scalability challenges as the number of species increases. This limitation arises because these methods use as input a large concordance factors (CFs) table, which summarizes the observed CFs of all possible four-species combinations in each row. The size of this table scales with the fourth power of the number of species, creating computational bottlenecks and highlighting the need for more efficient solutions. Sparse learning has been shown to reduce the dimensionality of large-scale datasets while producing results of comparable quality to those obtained using the full dataset. In this study, we adapted two sparse machine learning models—Elastic Net and Ensemble Learning + Elastic Net—to guide the subsample of an optimal number of rows from the CFs table required to accurately predict the overall phylogenetic network pseudolikelihood. Both methods account for the inherent correlation among rows, which arises because rows overlap in species information. We call this method *Qsin*. In two simulated datasets, Qsin reduced the dataset by approximately half without compromising accuracy. For the *Xiphophorus* fishes dataset, which contains 10,626 rows in the CFs table, we recovered the same topology as with the full CFs table but using only 763 rows. Using these subsamples also reduced running times by up to 60% without compromising accuracy. These gains are expected to persist as species numbers increase. Qsin contributes to ongoing efforts to make phylogenetic network inference more efficient and opens the door to analyses of more complex evolutionary histories. The source code for Qsin is freely available at: https://github.com/ulises-rosas/qsin.

## 1 Introduction

Phylogenetic networks are a generalization of phylogenetic trees (Kong et al., 2025). They model species evolution by accounting for signals from hybridization, gene flow, and reticulation (Solís-Lemus and Ané, 2016; Blischak et al., 2023). For instance, consider a phylogenetic network with a single hybridization event, resulting in two hybrid edges converging at a node. This scenario can produce two distinct underlying tree histories of species evolution, depending on which hybrid edge is followed. While both trees represent valid evolutionary histories, each may offer only a partial view of the species evolution. Phylogenetic networks, therefore, provide a more comprehensive representation of complex evolutionary histories.

One of the fastest algorithms for inferring phylogenetic networks is called SNaQ (Solís-Lemus and Ané, 2016). Currently, SNaQ models networks in which the sets of edges from hybridization cycles are all disjoint—known as level-1 networks. Among the reasons this method gained popularity includes that it i) accounts for incomplete lineage sorting (ILS), ii) models hybridization in small to medium-sized sets of taxa, and iii) does not rely on branch length information from gene trees, which can be sensitive to estimation error (Frankel and Ané, 2023). To reconstruct a phylogenetic network using SNaQ we typically start from the following table of concordance factors (CFs hereafter):

| 4-taxon set | $CF_1$ | $CF_2$ | $CF_3$ |
| --- | --- | --- | --- |
| $r_1$ | $f_{1,1}$ | $f_{1,2}$ | $f_{1,3}$ |
| $r_2$ | $f_{2,1}$ | $f_{2,2}$ | $f_{2,3}$ |
| ⋮ | ⋮ | ⋮ | ⋮ |
| $r_p$ | $f_{p,1}$ | $f_{p,2}$ | $f_{p,3}$ |

In the table above, let *r*_*j*_ denote a subset of four species in row *j* of the CFs table, obtained from the full set of species. Let *CF*_*k*_ represent topology *k* for this set of four species, also called a quartet topology. For example, for species A, B, C, and D, the three possible quartets are *AB*|*CD, AC*|*BD*, and *AD*|*BC*, where | indicates the edge separating sister species. Finally, f_*j,k*_ is the proportion of gene trees displaying quartet topology *k* for taxa in *r*_*j*_, hence the name concordance factors.

Given a proposed phylogenetic network topology *N*_*i*_, we calculate its log-pseudolikelihood score 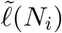 as:

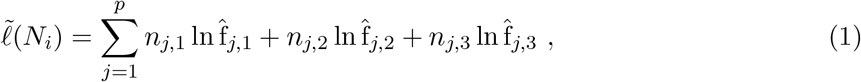

Where 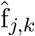 is the expected value of f_*j,k*_ obtained after branch length optimization, and *n*_*j,k*_ is the number of times quartet topology *k* for taxa in *r*_*j*_ appears among the observed gene trees. This score assumes that f_*j,k*_ follows a multinomial distribution and is used to search the network space: if the score increases for a new network topology, the topology is accepted; otherwise, the algorithm continues searching (Solís-Lemus and Ané, 2016). However, as the number of species increases, the SNaQ algorithm faces scalability challenges. This limitation stems from the quartic growth of the number of CFs rows with respect to the number of species. For example, with 50 species, SNaQ must evaluate the log-pseudolikelihood of all four-taxon sets, requiring the multiple assessments of 230,300 CFs rows per topology proposal. Moreover, as the number of species increases, the topology space expands rapidly—at least as fast as the tree space—further compounding the computational burden.

The challenge of working with large datasets is not exclusive to phylogenetics; it arises in many other fields as well (Yao and Wang, 2021). Typically, two scenarios are encountered: there are far more observations than variables (denoted as *n* ≫ *p*), or there are far more variables than observations (denoted as *p* ≫ *n*). One common approach to address such problems is through regression methods that impose sparsity constraints on the model weights—that is, forcing some weights to be exactly zero (Guyon and Elisseeff, 2003). For instance, to reconstruct a phylogenetic tree from a large DNA sequence alignment, Ecker et al. (2022) employed Lasso regression (Tibshirani, 1996). In their study, the variables were the site loglikelihoods calculated from random trees and the observations corresponded to the number of random trees used. Using Lasso regression, many variable weights were shrunk to exactly zero, and the authors used the non-zero weights, representing relevant sites to the overall loglikelihood, to reconstruct the tree. When the runtime of Lasso regression is lower than that of tree reconstruction, this approach is convenient because site loglikelihoods evaluations from random trees are inexpensive to compute: there is no need to optimize branch lengths, and one does not need to wait for the evaluation of a tree to finish before proposing the next one.

The above example shows how sparse regressions can effectively reframe the dimensionality reduction challenge posed by large datasets as a feature-selection problem (Lemhadri et al., 2021; Singh and Stufken, 2023). Intuitively, we could adapt a similar approach to improve the scalability of SNaQ. Instead of alignment sites, we would operate on the rows of the CF table. However, there are two key considerations we need to address: i) unlike sites in a sequence alignment, the rows in the CF table are not independent, and ii) in a high-dimensional settings (i.e., *p* ≫ *n*), a unique Lasso solution selects at most *n* features, where *n* is the number of observations (Rosset and Zhu, 2007; Hastie et al., 2015; Tibshirani and Wasserman, 2017), such as the chosen number of simulations in our case.

To address these issues, we propose the use of Elastic Net (Zou and Hastie, 2005), a sparse learning method that produces sparse solutions—also referred to as sparse learning. Elastic Net balances sparsity with the inclusion of correlated variables, enabling the selection of more than *n* non-zero coefficients when *p* ≫ *n*, by leveraging a grouping effect among correlated variables. Furthermore, we account for complex interactions among variables by modeling the log-pseudolikelihood score with a sparse ensemble of regression trees, also known as Importance Sampled Learning Ensemble (ISLE) (Friedman et al., 2003). Depending on the ensemble complexity, ISLE also has the added advantage of rapid testing of error convergence toward a specific sparse solution, allowing data-driven determination of the optimal percentage of row subsampling.

Here, we (1) propose an algorithm that reduces the number of rows in the CF table, guided by Elastic Net-based machine learning; (2) apply cross-validation (Hastie et al., 2009) to estimate all major hyperparameters of the machine learning models, including ISLE ensemble parameters; (3) analyze the time complexity of the algorithm as a function of the number of taxa; and (4) evaluate the algorithm on simulated data and on the best-characterized empirical network dataset for *Xiphophorus* fishes (Cui et al., 2013; Solís-Lemus and Ané, 2016). Our primary goal is to improve the runtime of phylogenetic network inference without compromising accuracy.

## 2 Materials and Methods

### 2.1 Regression Model Setup

Given a phylogenetic network *N*_*i*_, we define the quartet log-likelihood for row *r*_*j*_ in the CFs table as

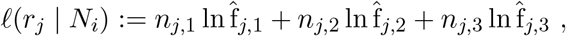

such that Eq. 1, which denotes the log-pseudolikelihood score, can be rewritten as

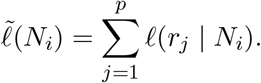

Similarly to Ecker et al. (2022), we then assume the existence of a scalar *β*_0_ ∈ ℝ and a vector *β* ∈ ℝ^*p*^ such that the calculation of 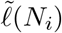 can be approximated by:

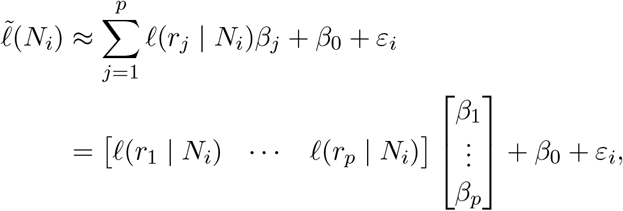

Where *ε*_*i*_ ∼ 𝒩 (0, *σ*^2^) is the error of the approximation with mean 0 and a constant variance *σ*^2^. For *n* phylogenetic networks, we obtain the following regression formulation:

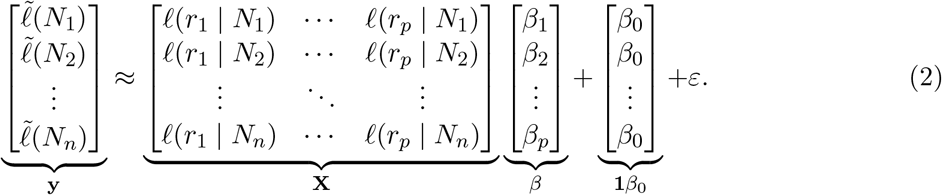

We define the left-hand-side response vector as **y** ∈ ℝ^*n*×1^, the predictor matrix as **X** ∈ ℝ^*n*×*p*^, and the right-hand-side vector as the intercept vector **1***β*_0_ ∈ ℝ^*n*×1^. A total of *n* phylogenetic networks can be obtained from simulations (see Section 2.2). Notice that there is a one-to-one correspondence between the columns of **X** and the rows of the CFs table. Specifically, column *j* of **X** represents the quartet log-likelihood of row *j* in the CFs table, given a set of *n* random phylogenetic networks. The coefficient vector *β* assigns weights to these columns.

If the weight for column *j* is set to zero (i.e., a sparse solution), its contribution to the overall prediction is negligible. Because of the direct correspondence between column *j* in **X** and row *j* in the CFs table, this implies that row *j* in the CFs table is also negligible and can be removed. Thus, the input size for SNaQ can be reduced in proportion to the sparsity of *β*. Using sparse learning techniques, as described in Sections 2.3 and 2.4, we obtain sparse versions of *β*.

### 2.2 Random Phylogenetic Networks

We conducted random phylogenetic network simulations using SiPhyNetworks (Justison et al., 2023). Following Kolbow et al. (2025), we used a speciation rate of 1, an extinction rate of 0.2, and drew inheritance probabilities from a Beta(10, 10) distribution. We also used a hybridization rate of 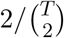 where *T* is the number of taxa. This hybridization rate causes hybridization events to occur more frequently toward the end of the process, when more lineages are available to choose from. As a result, more level-1 networks are generated, since the likelihood of hybridization events forming independent hybridization cycles increases. To accelerate lineage generation, we enabled only the lineage-generative mode. To ensure at least *n* level-1 networks, we simulated up to 4*n* networks and retained only those that satisfied the level-1 constraint. After filtering, we randomly permuted the tip labels and rescaled the branch lengths so that the average branch length was 1.25, approximately reflecting a moderate level of ILS (Kolbow et al., 2025).

Section 2.6 describes the simulated and empirical CFs tables used here. Following Eq. 1, we computed the quartet log-likelihood for each row in the CFs table across all *n* random phylogenetic networks generated using `topologyQPseudolik` in PhyloNetworks v0.16.4 (Solís-Lemus et al., 2017). In this study, we set *n* = 1000. Notably, this step is highly parallelizable, as each simulation—including the log-pseudolikelihood calculation—is independent.

### 2.3 Elastic Net

From the above data at Eq. 2 definition and after data centering, which allow us to omit the intercept term (Wang, 2022), we can define the following Elastic Net objective function:

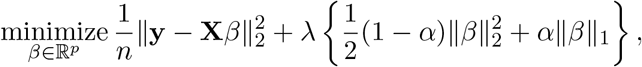

where *λ* is the sparsity parameter, and *α* is the weight between the *ℓ*_1_-norm and *ℓ*_2_-norm penalizations. The Elastic Net model is a generalization of the lasso model, and this becomes more apparent if we set *α* = 1. The addition of the *l*_2_-norm serves two purposes: it provides a grouping effect for correlated predictors, and it allows the selection of more than *n* solutions, a limitation of the lasso model (Rosset and Zhu, 2007). Hyperparameters *α* and *λ* are priors of the Elastic Net model and are usually obtained via cross-validation (CV) (Hastie et al., 2009, pp. 241).

Let Λ ∈ ℝ^*k*^ denote an ordered set of *k* values of *λ* arranged in decreasing order, and define the Elastic Net path as the set of *k* solutions corresponding to each *λ* ∈ Λ. Computing the full Elastic Net path is more computationally efficient than testing randomly selected *λ* values from the middle of the Λ range. This efficiency arises from the warm-start strategy: the solution obtained for a previous value, say *λ*_*k*−1_, serves as a good initial estimate for the model with the next value, *λ*_*k*_.

While the value of *α* is often pre-chosen on qualitative grounds (Hastie et al., 2009; Friedman et al., 2010), in this study we followed the 5-fold CV strategy proposed by Zou and Hastie (2005): from a relatively small set of candidate *α* values, each is used to compute the full Elastic Net path across all CV folds. The *α* that yields the lowest average CV error along its path is then selected.

The set we chose for *α* is {0.5, 0.95, 0.99}, and for the values of Λ, we let *k* = 100, where all the values are equally spaced on a logarithmic scale. The maximum value from this set, *λ*_max_ ∈ Λ, is the one that produces a fully sparse solution, and it can be theoretically obtained from data (Hastie et al., 2015, pp. 114). The minimum value from this set can be obtained as *λ*_min_ = *λ*_max_ · *ϵ*, where *ϵ* is a small constant. In this study, we set *ϵ* = 0.0001.

### 2.4 Importance Sampled Learning Ensemble (ISLE)

Applying Elastic Net directly to **X** constrains the prediction of **y** to a linear function of the predictor variables, which can lead to underfitting. To address this limitation, we relax the linearity assumption, allowing arbitrarily nonlinear functions of the predictor variables in predicting **y**. This, in turn, allows non-parametric interactions among subsets of rows from the CFs table to predict the log-pseudolikelihood score of a phylogenetic network, while still retaining the sparsity benefits of Elastic Net. We achieve this using the ISLE algorithm (Friedman et al., 2003; Seni and Elder, 2010; Zeng, 2022), which consists of two main steps: (i) generating a strong ensemble of nonlinear basis functions, and (ii) post-processing of these functions with a sparse regression method such as Elastic Net to select a subset with optimal predictive performance. In this study, we use regression trees as the nonlinear basis functions.

Let the prediction of *n* log-pseudolikelihood scores from the matrix **X**, as defined in Eq. 2, using a regression tree be denoted as 𝒯 (**X**; *γ*_*j*_) ∈ ℝ^*n*×1^, where *γ*_*j*_ is the optimal set of parameters in the regression tree (e.g., splitting variables at internal nodes, tree depth). Then, we can form the following matrix **T** ∈ ℝ^*n*×*M*^ containing the predictions of *n* log-pseudolikelihood scores from *M* regression trees:

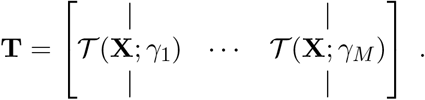

Let **w** ∈ ℝ^*M*^ be a vector containing a weight for the prediction performance of each regression tree; then, by making this vector **w** sparse using Elastic Net on the centered data, we can create a subsample of these regression trees:

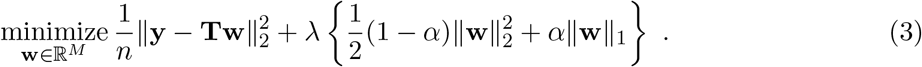

However, if all regression trees are generated randomly, we may undersample important variables—in our case, informative rows of the CFs. Then, the ISLE algorithm also proposes leveraging an importance sampling approach to obtain a stronger ensemble (Seni and Elder, 2010, pp. 43). The details of the algorithm in the specific case of regression trees can be found in Algorithm 1. *S*_*m*_(*η*) represents the *m* subsample of *n* · *η* observations, where *η* ∈ (0, 1] is a constant. *ν* ∈ [0, 1] is another constant that represent the memory rate. Intuitively, if we decrease *η*, then we increase the randomness of the regression trees; and if we increase *ν*, then the regression trees avoid similarity between them. Friedman et al. (2003) recommended using *ν* = 0.1 and *η* ≤ 0.5 for very shallow regression trees.

Via 5-fold CV, we tested the hyperparameter values near recommended defaults: five equally spaced *η* values from 0.0316 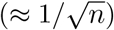 to 0.5, five equally spaced *ν* values from 0.01 to 0.1, and maximum leaf nodes of 2, 3, 4, and 5. More generally, as the number of possible combinations of *η, ν*, and leaf node counts increases, we randomly selected up to 100 hyperparameter combinations, following the strategy of Bergstra and Bengio (2012).

Each of these 100 ISLE ensemble hyperparameters was paired with the three values of *α* introduced in Section 2.3 for the Elastic Net path, yielding a total of 300 tested hyperparameters (100 × 3). All *M* regression trees were constructed using scikit-learn v1.5.1 (Pedregosa et al., 2011). In this study, we set *M* = 4000.

### 2.5 Phylogenetic Network Inference

Once we obtain a sparse model—either from Elastic Net or ISLE, as described in the previous sections—we aim to identify the variables in **X** that have been selected. These selected variables correspond to specific rows in the CFs table, enabling the inference of a phylogenetic network from this reduced table.

In the case of Elastic Net, the selected variables are those associated with non-zero coefficients in the solution vector *β*, given a particular pair (*λ, α*) from the Elastic Net path. Specifically, let

#### Algorithm 1

Ensemble generation for the ISLE algorithm with regression trees as base learners (Friedman et al., 2003; Seni and Elder, 2010)

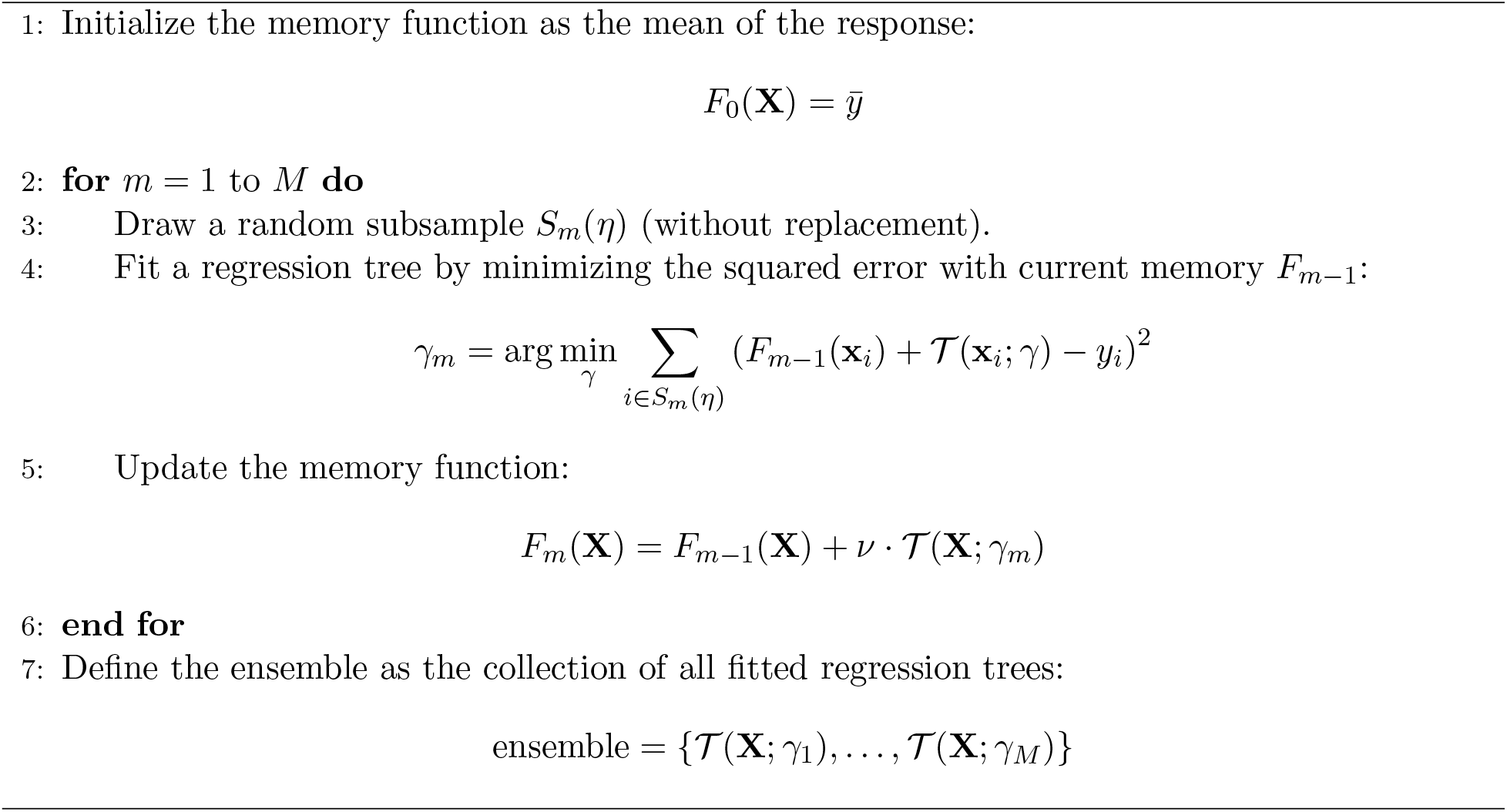

*λ*_*k*_ ∈ Λ denote the *k*^th^ value of *λ* tested, and let 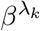 be the corresponding solution. The index set 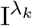 representing the selected variables, is defined as:

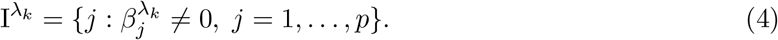

A similar process applies with ISLE (see Algorithm 2). Here, we examine the splitting variables used by the regression trees selected in the ISLE ensemble instead of non-zero coefficients in *β*. Selection is again determined via the Elastic Net path. Let 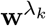 denote the ISLE solution for a given *λ*_*k*_ ∈ Λ and *α*, where 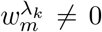 indicates an active regression tree *m* in the ensemble. Let *X*_*j*_ ∈ *γ*_*m*_ denote a splitting variable *j* used in the set of optimal parameters *γ*_*m*_ that define regression tree *m*. The index set 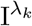 is then constructed as described in Algorithm 2. Importantly, the index set 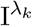 must always include variables that collectively cover all species.

We now summarize all previously described steps in Algorithm 3, which we refer to as *Qsin* (Quartet Subsampling for Inference of Networks). Notably, subsampling guided by Elastic Net instead of ISLE can also be viewed as part of Qsin by skipping step 2, setting **T** = **X** in step 3, and using the row selection defined in Eq. 4 instead of Algorithm 2 in step 4. We refer to this variant as Qsin without ensemble creation. For phylogenetic network inference using SNaQ and subsamples (step 5 in Qsin), we used the default parameters and performed 15 independent runs, unless otherwise specified.

Detailed analyses of the time complexity for each step in Algorithm 3 are provided in Supplementary Material S.II. Building upon these individual results, we also derive the overall computational complexity of the complete algorithm, which we formalize in the following Theorem 1:

#### Theorem 1

(Proof available in Supplementary Material S.II). *Let T be the number of taxa, n be the number of simulations, M the number of used regression trees, and ρ* ∈ (0, 1] *the fraction of*

#### Algorithm 2

Compute the index set of CFs rows from a subsample of regression trees.

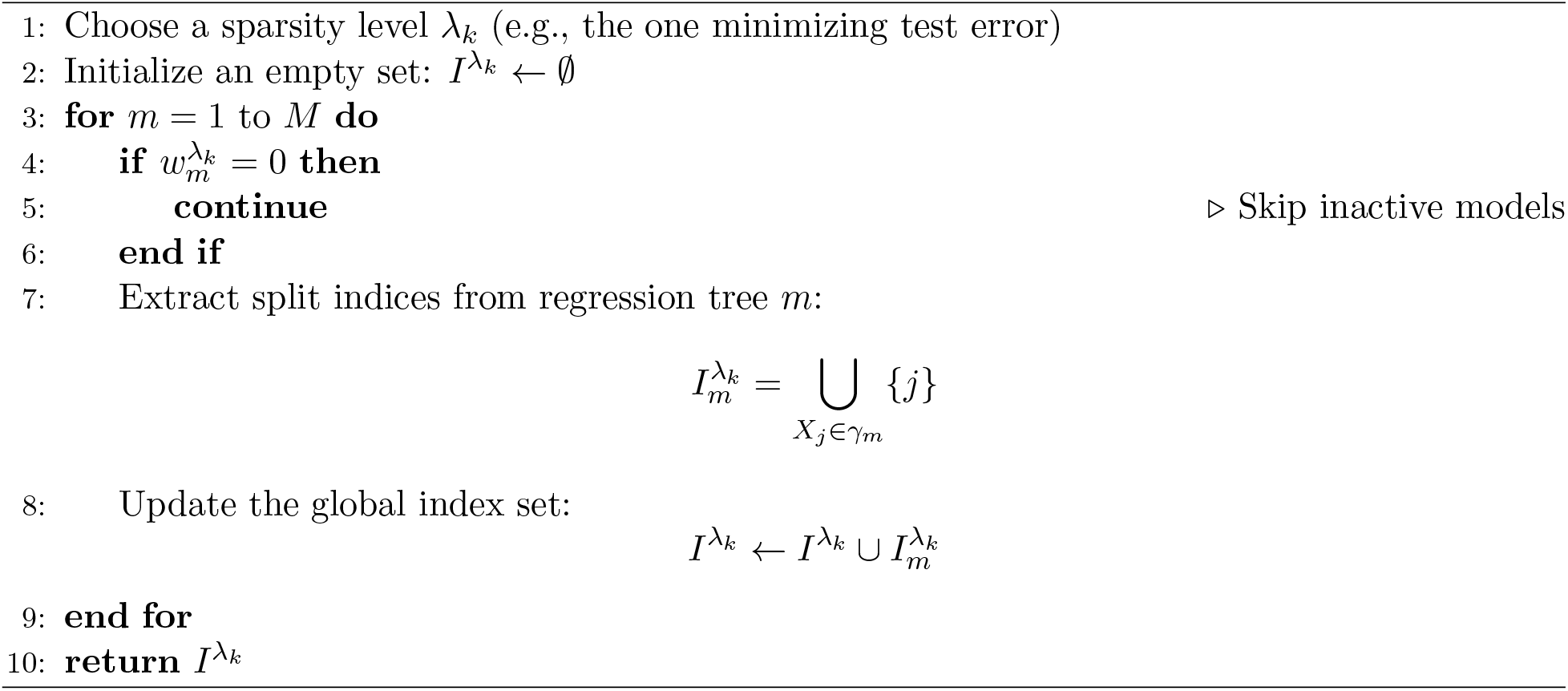

*selected CFs rows used in reconstruction. Assume the number of SNaQ iterations is at most on the order of T*. *Then, the total time complexity of Algorithm 3 is O nT* ^5^ + *MT* ^4^*n* log *n* + *ρT* ^7^.

We can also consider two extreme cases for *ρ*: one where all rows are selected and another where only enough rows are chosen to cover all species. By instantiating *ρ* under these two extreme scenarios, we can define the boundaries of the speed-up, within which the observed speed-up can lie:

#### Corollary 1

(Proof available in Supplementary Material S.III). *Let T be the number of taxa. Under the same assumptions outlined in Theorem 1, and as T* → ∞, *the worst-case scenario yields a speed-up factor of O*(1) *(i*.*e*., *no speed-up) and the best-case scenario yields a speed-up factor of O*(*T*^3^) *relative to SNaQ*.

Therefore, as the number of species increases and *ρ* falls below 1, a speed-up in runtime can generally be expected.

#### Algorithm 3

Qsin, a pipeline to reconstruct phylogenetic networks.

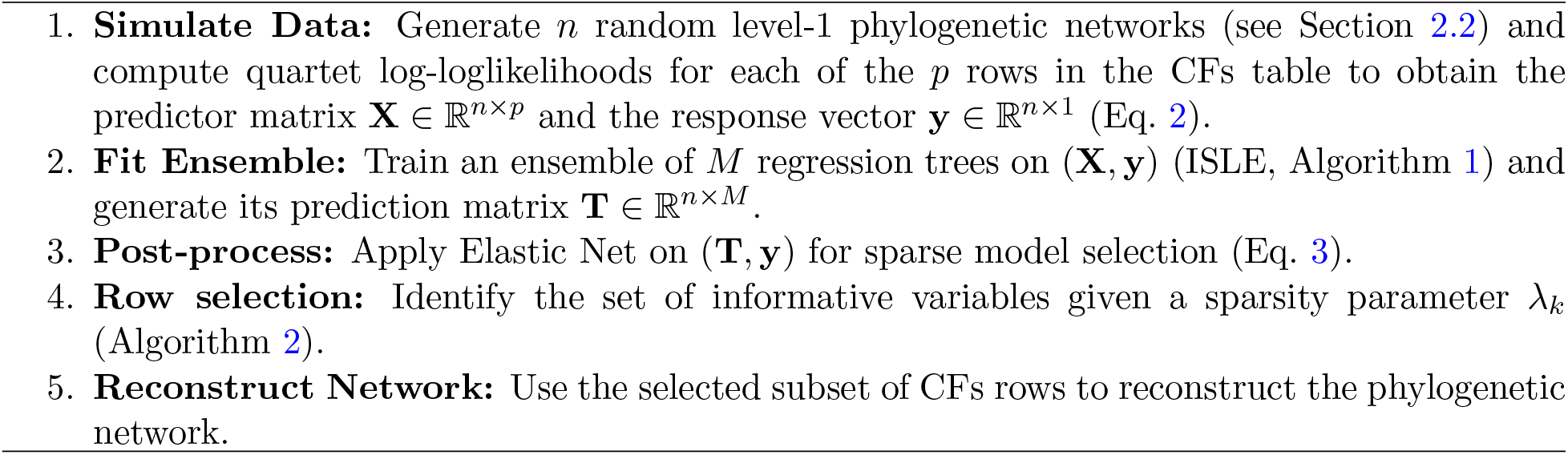

### 2.6 Model Evaluation

We used datasets presented in Solís-Lemus and Ané (2016) to evaluate the performance of the proposed pipeline. Two simulated datasets were included: the first contains 6 species, resulting in 15 rows in the CFs table based on 100 gene trees, and includes one hybridization event. The second simulated dataset contains 15 species, resulting in 1,365 rows in the CFs table based on 300 gene trees, and includes three hybridization events. A third dataset consists of empirical data from *Xiphophorus*, comprising 24 species and 10,626 rows in the CFs table based on 1,183 gene trees. Although the true number of hybridization events in *Xiphophorus* is unknown, we set *h*_*max*_=2 in SNaQ, as this value was found to be optimal using the slope heuristic in Solís-Lemus and Ané (2016).

The quality of log-pseudolikelihood predictions was assessed using the root mean squared error (RMSE), defined as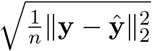 where **ŷ** denotes the predicted values of **y** (the observed log-pseudolikelihood score; see step 1 in Algorithm 3). For simulated datasets, where the true network is known, we evaluated topological accuracy using the hardwired distance between the inferred and true networks. This metric, implemented as `hardwiredclusterdistance` in PhyloNetworks v0.16.4 (Solís-Lemus et al., 2017), captures the topological dissimilarity between two networks. Additionally, we computed the distance to the true network while ignoring the direction of the hybrid edge, referred to as the unrooted distance. To do this, we shifted the position of the hybrid node within the hybridization cycle of the inferred network and recalculated the distance. The new position of the hybrid node was chosen so that its descendant taxa matched those of the hybrid node in the true network. A custom script, named `ChangeDirection.py`, for performing this network manipulation is available in the accompanying GitHub repository: https://github.com/ulises-rosas/qsin.

We applied this unrooted distance to the second simulated dataset, which included 15 species, because it contains a hybridization cycle with four nodes and few taxa attached to the corresponding subnetworks, making the inference particularly challenging (see Fig. 9 in Solís-Lemus and Ané (2016)). As noted in that study, the unrooted distance provides a more appropriate measure of performance for this dataset. Accordingly, all comparisons for this dataset were made using the unrooted distance.

## 3 Results

When applying Elastic Net directly to the first simulated dataset (i.e., Qsin algorithm without ensemble creation), which included 15 rows in the CFs table, we found that the Elastic Net path selected more rows as the sparsity parameter decreased (Fig. 1). In the Elastic Net path (Fig. 1A), we subsampled across the region between the point at which no rows are selected (i.e., the maximum sparsity parameter is used) and the point at which all rows are selected. We found that a subsample consisting of approximately half of the dataset achieved an accuracy of 93% (Fig. 2A), as measured by the proportion of times the inferred network had zero distance from the true network. This accuracy was the same as when using the complete dataset (93%, Fig. S.2A). Additionally, reducing the number of rows also decreased runtime for phylogenetic network inference (Fig. 2B): while processing the full dataset required a median of 47.19 seconds, the subsampled version completed in a median of 39.77 seconds. Obtaining the subsample itself required 28.98 seconds. There was approximately a 15% decrease in network inference running time on this small dataset.

**Figure 1.**
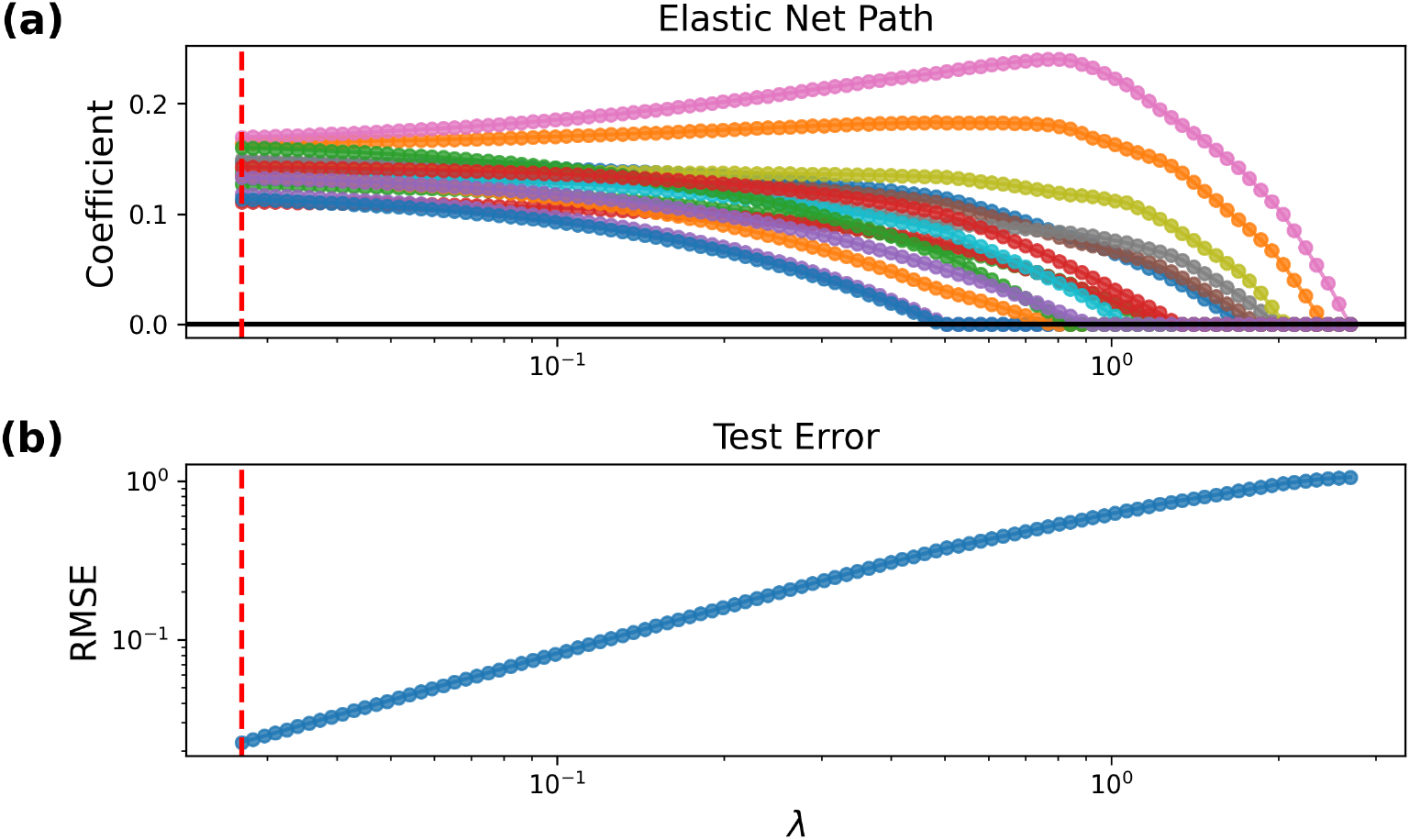
The Elastic Net path was directly applied to a CFs table with 15 rows, involving 6 species. Elastic Net alone tended to keep adding rows without a clear point of convergence, as test error steadily decreased until nearly all rows were included. In **(a)**, each colored line represents a row in the CFs table. From right to left, we observe that when *λ* is at its largest value, no rows are selected—as indicated by all colored lines being at zero. Dashed red vertical line indicates the point in the path with the lowest error, which is when all the rows are active in the prediction. **(b)** shows the testing RMSE along the path, which decreases almost monotonically as more rows are selected. Before reaching the lowest point, the testing error does not show sign of converging. The best *α* obtained via 5-fold CV was 0.5.

**Figure 2.**
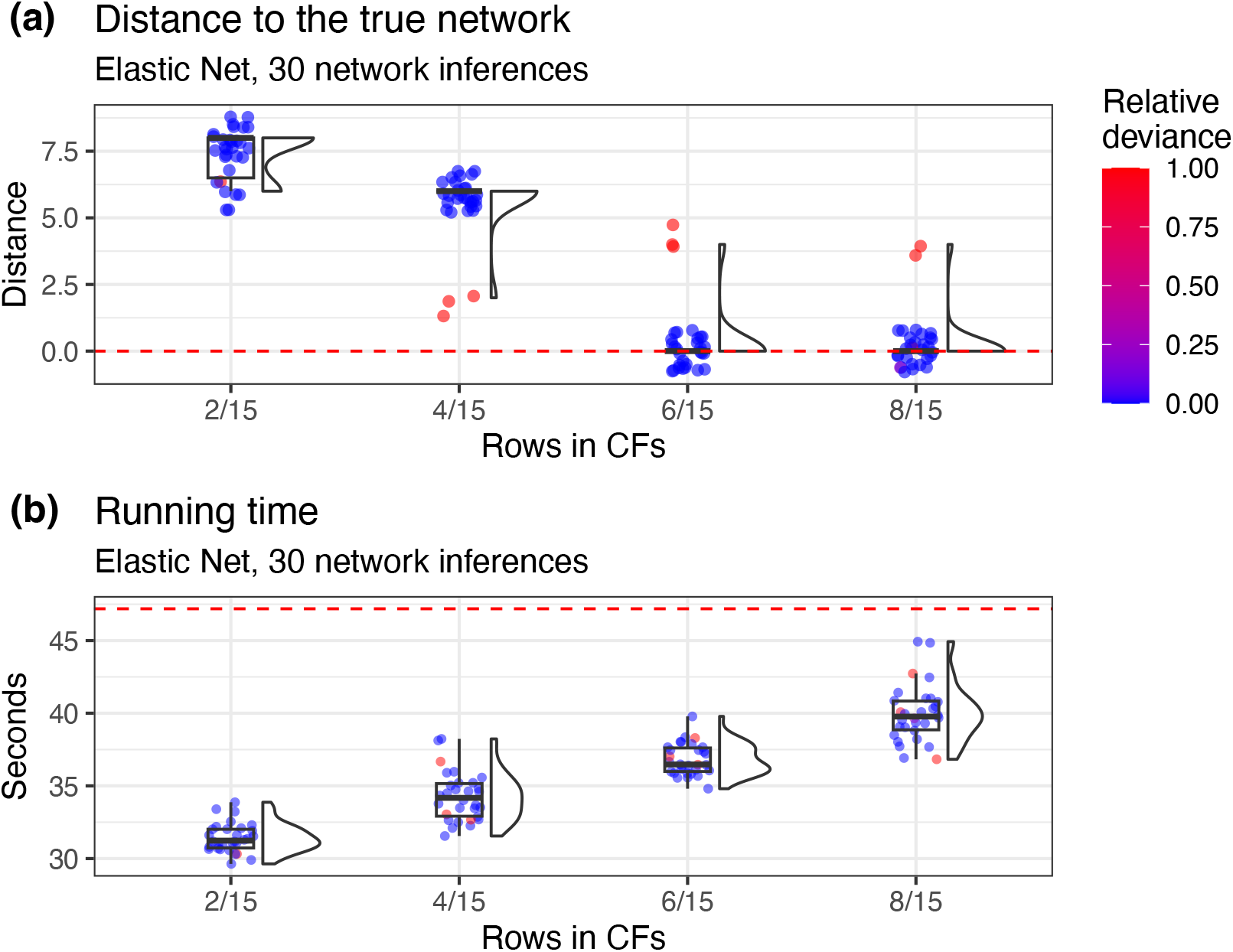
Accuracy of reconstruction using subsampling guided by Elastic Net, i.e, Qsin algorithm without ensemble creation, on a CFs table with 15 rows. **(a)** shows the distance to the true network. With 8 rows, the true network is recovered 93% of the 30 repetitions. The red dashed line indicates zero distance. **(b)** shows inference time for different row selection sizes. The red dashed line indicates the median time required to reconstruct the network using the full dataset, which is approximately 47.2 seconds. Each repetition used 10 independent runs. Random selection of 8 rows had lower accuracy (Fig. S.6A). Relative deviance denotes the normalized log-pseudolikelihood score. In summary, results show that using Elastic Net to select about half of the CF rows recovered the true network in most runs while reducing inference runtime by approximately 15%, whereas random subsampling of the same size failed.

We observed similar performance when we modified step 1 of Qsin algorithm (see Algorithm 3) by drawing the branch lengths of the random networks from an exponential distribution (mean = 1.25; Fig. S.3) or left unmodified (Fig. S.4). However, when the branch lengths were optimized to obtain log-pseudolikelihood scores using `topologyMaxQPseudolik` instead of `topologyQPseudolik` (see Section 2.2) in PhyloNetworks v0.16.4, a subsample of half the dataset resulted in 0% accuracy (Fig. S.5). Randomly selecting rows of the same size as in Fig. 2 also yielded 0% accuracy (Fig. S.6A).

While Elastic Net can successfully generate optimal subsamples on the first simulated dataset, there was no clear point along the path at which to stop subsampling, as the testing error did not converge to a distinct minimum within the region between no row selection and full row selection (Fig. 1B). Therefore, we proceeded with the ensemble learning-based approach of Qsin (see Algorithm 3).

The ensemble of trees used by Qsin in step 2 introduces additional hyperparameters. To visualize this hyperparameter space, we used the second simulated dataset, holding out a random fold of the data and fixing *α* = 0.5 (Fig. 3). We observed that varying the level of nonlinearity in the sparse ensemble—measured by the maximum allowed number of leaves in each regression tree—resulted in different prediction errors for the log-pseudolikelihood score. Across all tested values for the number of leaves, prediction error increased as *η* approached 0.5, a region corresponding to fewer random ensembles. The optimal value of *ν*, the hyperparameter controlling ensemble diversity, varied depending on the number of leaves. We also observed that increasing the number of leaves generally reduced the prediction error. For ensembles with a maximum of 2 and 5 leaves, the lowest errors were 0.175 and 0.155, respectively, in this held-out fold. However, increasing the number of leaves also led to less sparse solutions. For example, the best solution with 5 leaves included 1,258 out of 1,365 rows in the CFs table (92.2%).

**Figure 3.**
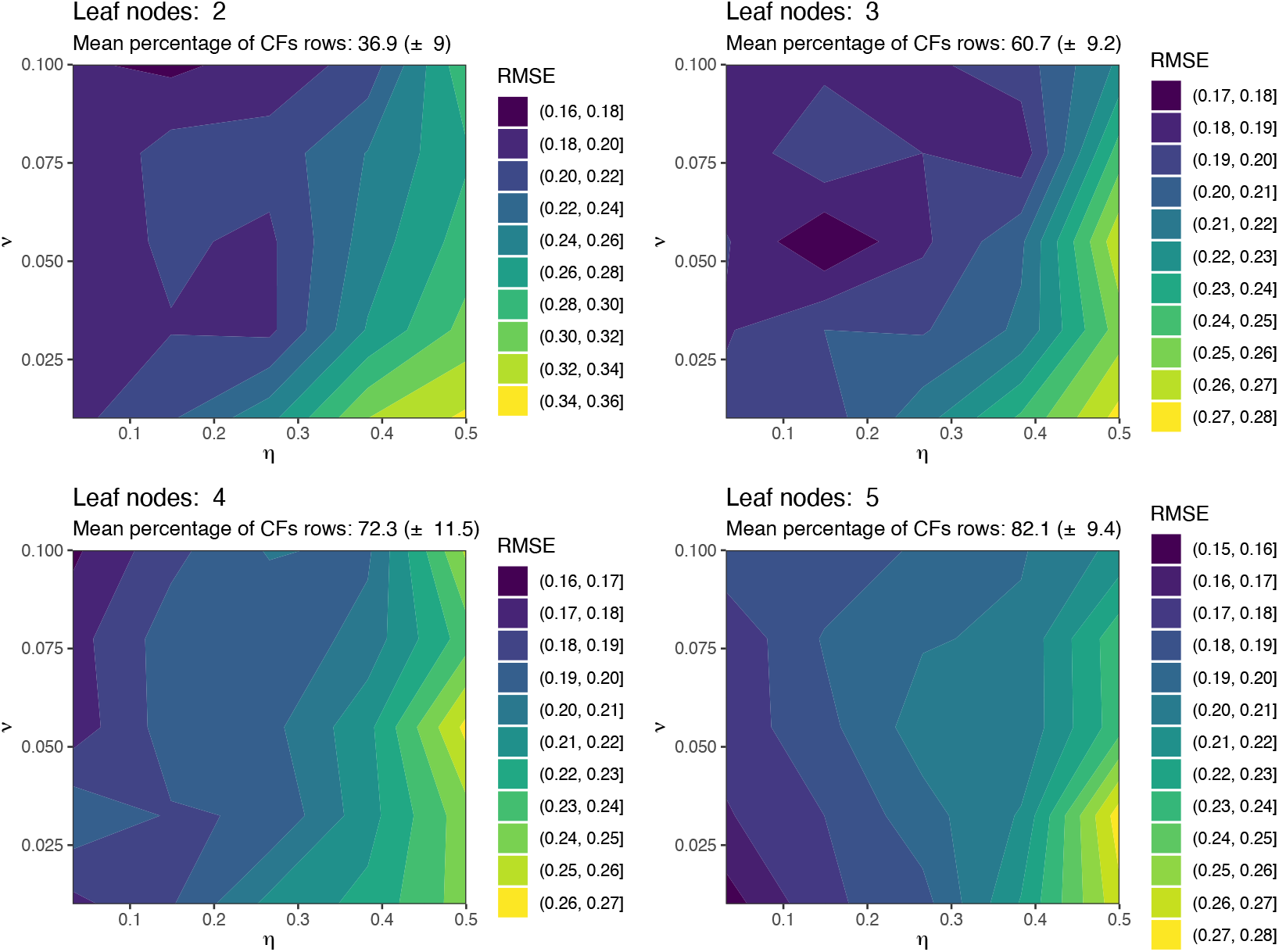
Hyperparameter space of ISLE ensemble with different level of non-linearity as measured by the number of leaf nodes in the regression trees. The darkest area in each panel represents the space with the lowest testing error, RMSE. The grid exploration is based on one realization of the training and testing set with random assignment. All the RMSEs represent the lowest error in the Elastic Net path. In summary, ensembles with 2 leaves achieved the best balance of accuracy and sparsity, while 5-leaf trees selected most of the rows without a clear gain in sparsity.

Since our goal is to obtain highly sparse solutions, we fixed the maximum number of leaves to 2 and performed a 5-fold CV to estimate the remaining hyperparameters: *α, η*, and *ν*. Across the five folds, the hyperparameters that minimized the average prediction error were *α* = 0.5, *η* = 0.0316, and *ν* = 0.1. Applying the Elastic Net path to the ensemble generated with these hyperparameters, we observed that the testing error reached its minimum between the extremes of no row selection and full row selection (Fig. 4). At the point of minimal error (0.178), as measure by RMSE along the path, 750 regression trees were selected, which, via Algorithm 2, spanned 578 rows of the CFs table (42.3%).

**Figure 4.**
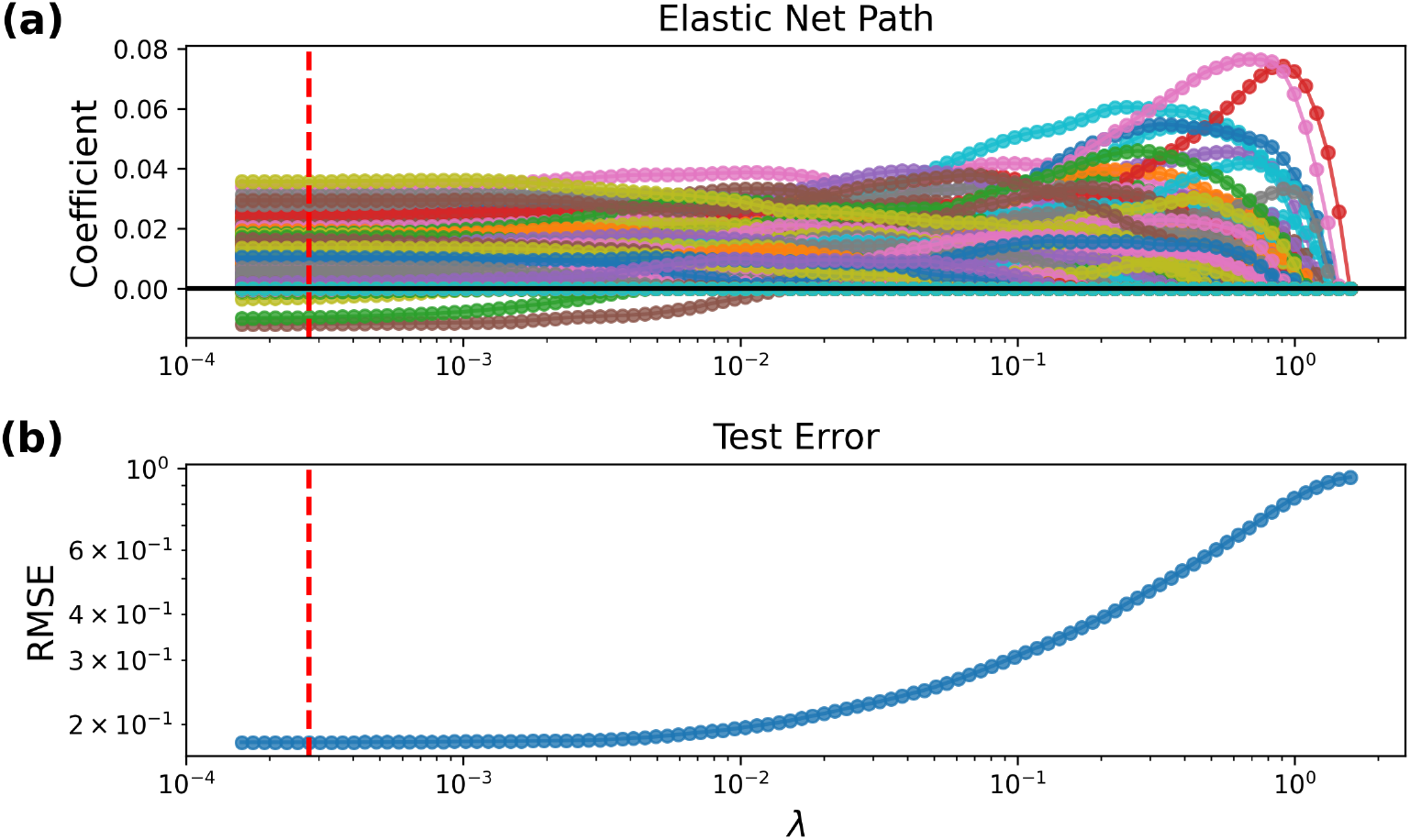
The Elastic Net path applied to the ISLE ensemble on a CFs table with 1,365 rows, involving 15 species. Hyperparameters were set to 2 leaf nodes, *η* = 0.0316, and *ν* = 0.1. In **(a)**, each colored line represents a regression tree. From right to left, we observe that when *λ* is at its largest value, no regression trees are selected — as indicated by all colored lines remaining at zero. The red dashed vertical line marks the point along the path with testing error convergence. **(b)** shows the testing RMSE along the elastic net path, reaching its testing error convergence when 750 out of 4,000 regression trees are active—corresponding to 578 rows in the CFs table. In summary, using the ISLE ensemble, prediction error converged well before all trees were active, so that only a fraction of the rows (42.3%) was enough to reach stable accuracy.

As in Fig. 1, we created equally spaced subsamples along the Elastic Net path, from close to the beginning up to the point of minimal error (i.e., optimal subsample), and reconstructed phylogenetic networks using only these row subsamples (Fig. 5). We observed a smooth increase in accuracy as the subsample size approached the optimal subsample, which achieved an accuracy of 70%. This accuracy was essentially the same as that obtained using the complete dataset (69%; Fig. S.2B). In contrast, random selection of the same number of rows resulted in a lower accuracy of 58% (Fig. S.6B). Although both subsampling strategies had a median unrooted distance to the true network of 0, the interquartile range of these distances was 2 for the optimal subsample and 10.5 for random selection. Thus, the optimal subsample not only improved accuracy but also produced networks with less variability. Additionally, reducing the number of rows also decreased runtime for phylogenetic network inference: while processing the full dataset required a median of 10.76 hours, the subsampled version completed in a median of 4.25 hours. Obtaining the subsample itself required only 0.71 hours. There was approximately a 60% decrease in network inference running time.

**Figure 5.**
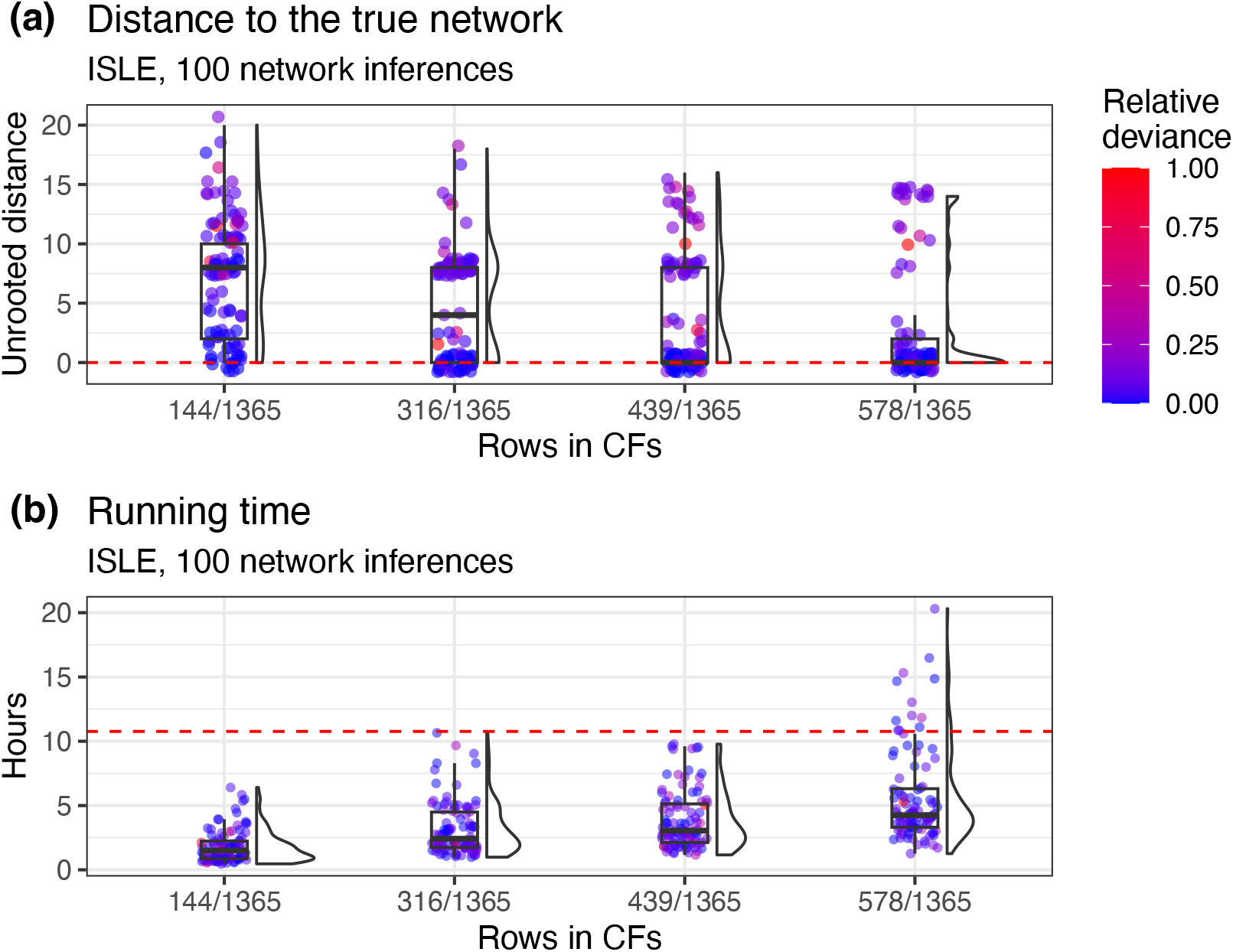
Accuracy of reconstruction using subsampling guided by ISLE, i.e., Qsin algorithm, on a CFs table with 1365 rows. **(a)** shows the unrooted distance to the true network. With 578 rows, the unrooted true network is recovered 70% of the 100 repetitions. The red dashed line indicates zero distance. **(b)** shows inference time for different row selection sizes. The red dashed line indicates the median time required to reconstruct the network using the full dataset, which is approximately 10.76 hours. Random selection of 578 rows had lower accuracy (Fig. S.6B). Relative deviance denotes the normalized log-pseudolikelihood score.. In summary, ISLE-guided subsampling reliably recovered the true network in most runs while reducing runtime by ≈ 60%, whereas random subsampling of the same size did not achieve similar accuracy.

The speed-up behavior shown in Fig. 2 and Fig. 5 is consistent with the theoretical result stated in Corollary 1. As the number of species increases and the fraction of selected rows in the CFs table, denoted by *ρ*, is less than 1, we can generally expect a speed-up in runtime.

We next applied Qsin, as described above, to the *Xiphophorus* dataset. Fixing the maximum number of leaves to 2, the optimal set of hyperparameters obtained via 5-fold CV was *α* = 0.5, *η* = 0.1487, and *ν* = 0.0775. The Elastic Net path generated from this configuration resembled that observed in the simulated dataset: the prediction error converged before reaching its minimum, at which point 763 out of 10,626 rows from the CFs table (7.81% of all rows) were selected (Fig. S.7). Since there was variability in inference accuracy for the previous datasets, we repeated the inference six times using different random seeds and selected the two networks with the highest log-pseudolikelihood scores, using the 7.81% subsample and *h*_max_ = 2. The network with the highest score of −709.19 matched exactly the most supported network topology described in Solís-Lemus and Ané (2016), with *h*_max_ = 2 (Fig. 6). Similarly, the second-best network (−742.63; Fig. 6B) matched the second most supported topology in that study (Fig. 6C). All remaining networks with lower scores captured different features of the top-scoring networks. For example, the first hybridization event originating from the branch of *X. xiphidium* to the branch of the most recent common ancestor of *X. cortezi* and *X. continens* (Fig. S.8A, B, C); others matched the second most supported network shown in Fig. 6B (Fig. S.8C), or matched the major tree—defined as the tree obtained by removing all hybrid edges with inheritance probabilities below 0.5—which corresponds to the major tree reported in Solís-Lemus and Ané (2016) for both *h*_max_ = 0 and *h*_max_ = 1 (Fig. S.8A, D). Finally, we increased the number of independent runs from 15 to 25 and then to 30. In these extended runs, a single repetition recovered the top-scoring network (Fig. S.9). In contrast, randomly selected rows of the same size did not recover the top-scoring network in any of these extended runs (Fig. S.10).

**Figure 6.**
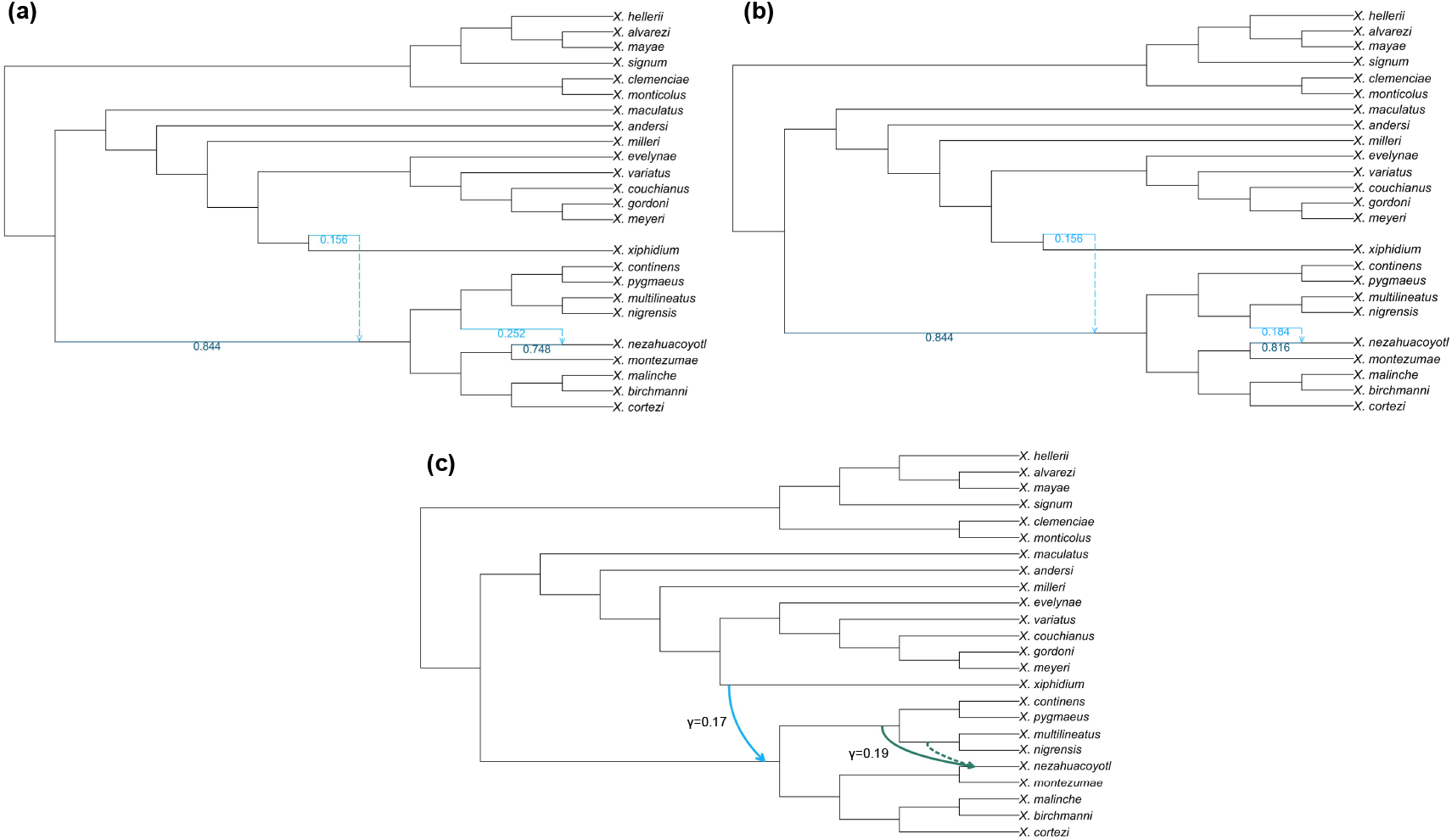
Phylogenetic network reconstruction for the *Xiphophorus* dataset using a subsample of 763 rows (7.81% of the CFs table) and the two best log-pseudolikelihood scores. **a)** shows the network with a log-pseudolikelihood score of −709.218, and **b)** shows the one with −742.635. The subsample was obtained using the ISLE algorithm with hyperparameters set to *α* = 0.5, two leaf nodes, *η* = 0.1487, and *ν* = 0.0775. The hyperparameters *α, η*, and *ν* were selected via 5-fold CV. For reference, **c)** shows the best networks obtained in Solís-Lemus and Ané (2016). The solid lines indicate the most supported network, while the skyblue solid line and green dashed line represent the second most supported one.

## 4 Discussion

We demonstrate that redundancy in the CFs table can be effectively exploited through machine learning–guided subsampling (Qsin, Algorithm 3). This subsampling approach accounts for the non-independence among CFs rows and can greatly reduce running times. In two simulated datasets, we were able to recover the true networks with high accuracy using only about half of the data. Such reductions yielded results comparable to those obtained with the complete dataset. In the empirical dataset, we found that the same phylogenetic network could be reconstructed as when using the entire dataset. The reduction in running time achieved by decreasing the number of rows is consistent with the theoretical result stated in Theorem 1 (Corollary 1).

Sparse learning has shown great promise for reducing model complexity across domains, including neural networks (Srivastava et al., 2014; Lemhadri et al., 2021). The core principle is that, even if the true data-generating process is unknown, it can often be represented in a lower-dimensional space, allowing variance and noise to be reduced (Hastie et al., 2009, pp. 610). Previous approaches using sparse learning for data size reduction first fixed an expected subsample size and then selected a sparse model that best matched that number of variables (Ecker et al., 2022). In contrast, our approach fixed model complexity, as measured by the number of leaf nodes, and allowed the data to determine the optimal subsample size. To our knowledge, this represents the first use of a non-parametric sparse model such as ISLE for data reduction in phylogenetics. This reduction is possible because the ISLE algorithm typically converges quickly to a solution along the Elastic Net path (Hastie et al., 2009, pp. 620), owing to its importance sampling strategy (Seni and Elder, 2010). This strategy favors regression trees that resample variables most relevant for improving prediction (i.e., building a strong ensemble) (Liu et al., 2022). Adding variables outside this set of important predictors tends only to introduce noise into the ensemble, or at best, provide no improvement—a behavior that is consistent with our ISLE solutions (Fig. 4, Fig. S.7).

A non-parametric model such as ISLE appears necessary because relationships among quartet log-likelihoods of rows in the CFs table can be far more complex than linear, particularly when predicting the overall log-pseudolikelihood score in the presence of beyond-level-1 phylogenetic networks, an area of active research (Allman et al., 2025a). Since our goal was to achieve substantial data reduction, we restricted the ensemble to trees with only two leaf nodes, also known as stump trees (Oliver and Hand, 1994). Although these regression trees are very simple, they can still yield complex relationships when used in an ensemble (Seni and Elder, 2010). For instance, they were the first trees used in AdaBoost algorithms (Freund et al., 1996) and have more recently been applied to improve out-of-distribution predictions in deep learning (Andriushchenko and Hein, 2019).

The Qsin algorithm makes use of Elastic Net to guide the subselection which is more suitable in high dimensional problems (*p* ≫ *n*) than Lasso. Unlike Lasso, the number of variables selected by Elastic Net is not upper-bounded by the number of observations (*n*) (Friedman et al., 2010; Gaines et al., 2018), and it also accommodates correlated variables by leveraging a grouping effect. When variables are correlated, Lasso tends to pick one and ignore the rest, leading to have erratic behavior in the path (Friedman et al., 2010). Accounting for variable correlation is important as rows in the CFs table are inherently not independent. In the context of phylogenetics, this is not the first study to recognize the importance of correlation when implementing sparse learning methods. For instance, in the *ℓ*1*ou* method for selecting the optimal number of shifts in trait optima, Khabbazian et al. (2016) accounted for the correlation of the coefficients in their sparse learning model using Lasso with a backward selection step that discards variables not improving the information criterion. More recently, the ESL method for associating trait presence or absence with a hot-encoded DNA alignment, Kumar and Sharma (2021) accounted for the cluster in groups of different genes by using grouped Lasso. Thus, when strong correlations among variables are expected, sparse learning models that explicitly address these correlations are preferable.

Random selection had lower performance across all tested datasets compared to Qsin selection. It also showed greater variability in its unrooted distance to the true network in the second simulated dataset with 15 species. However, in this same dataset, random selection of rows achieved an accuracy of about 58%, outperforming naïve random inference of the phylogenetic network. The utility of random subsampling in large datasets has also been observed in phylogenetic tree inference from quartets and is widely employed in methods such as SVDquartets (Chifman and Kubatko, 2014). This effectiveness further reinforces the idea that the dimensionality required to evaluate the fit of a phylogenetic hypothesis (e.g., number of sites in the alignment, number of gene trees, number of rows in the CFs) lies in a much lower-dimensional space. Consequently, we can exploit this redundancy using statistically principled algorithms (Vachaspati and Warnow, 2015; Warnow, 2018; Gupta et al., 2025), as demonstrated here.

Previous work has demonstrated the advantages of sparse learning for improving runtime (Khab-bazian et al., 2016; Ecker et al., 2022), and our study confirms these benefits. Beyond empirical evidence, we also theoretically show that reductions in running time necessarily occur whenever a subsample of the dataset is taken (Corollary 1) and as the number of taxa *T* approaches infinity. These derivations align with intuition—the less data, the fewer computations—but our analysis further provides a precise characterization of how these savings scale with *T*, *n, M*, and *ρ*, thereby clarifying the conditions under which subsampling yields the greatest gains. As the number of species increases, the computational benefit described in Eq. 1 is primarily influenced by the fraction of rows selected, *ρ*. While we can set *ρ* = 1*/T* to consistently achieve an order-of-magnitude speed-up compared with conventional inference, *ρ* is ultimately data-driven, and forcing it otherwise may not yield optimal solutions. Nonetheless, the Qsin algorithm effectively addresses the data size problem, which represents the first step in inference, and can also serve as a preprocessing stage for recent algorithmic advances such as divide-and-conquer approaches (Kolbow et al., 2025; Allman et al., 2025b) in phylogenetic network inference.

One limitation of the Qsin algorithm is the influence of parameters such as *n* and *M* on the overall time complexity (see Theorem 1). These parameters affect the runtime of steps such as log-pseudolikelihood evaluation, *O*(*nT* ^5^), and the regression step, *O*(*MT* ^4^*n* log *n*) (see Supplementary Material S.II)—both of which remain less costly than network inference when *T* is sufficiently large. However, these parameters can be reduced, forcing the sparse learning model to search for the best solution within a more constrained space, thereby yielding overall runtime improvements even for datasets with relatively few taxa. Another limitation is that the sparse learning model presented here does not incorporate network topology information. Instead, we rely on the rationale that an accurate prediction of the log-pseudolikelihood score implies an accurate prediction of the network topology. A more direct approach should consider both simultaneously. For example, Voznica et al. (2022) and Xie et al. (2025) successfully encoded phylogenetic trees in their machine learning models. However, the most effective data representation of phylogenetic network topology for machine learning models—particularly sparse models—remains an open question. Future improvements should also consider subsampling not only the full regression trees, as we did here, but potentially their components (e.g., internal nodes), an approach recently shown to improve ensemble accuracy (Liu et al., 2025).

On the other hand, if we want greater control over the frequency of taxa selected—in addition to the number of rows—we can use the constrained Lasso (Gaines et al., 2018), a direction that remains underexplored in phylogenetics. For example, let **z**_*t*_ ∈ ℝ^*M*×1^ be the column vector encoding the presence or absence of taxon *t* in an ensemble of *M* regression trees, such that each element in the vector is

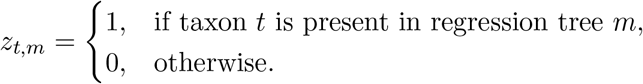

If we constrain **w** in Eq. 3 to take values from 0 to 1, then **w** can be interpreted as the ensemble posterior probabilities (Hastie et al., 2009, pp. 290). The weighted sum 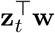 approximates the weighted average frequency of taxon *t* across the ensemble subselection. We can further require this weighted sum to be at least some constant, thereby enforcing a minimum frequency for each taxon (see Supplementary Material S.V for the incorporation of these constraints into Eq. 3). This approach offers a promising direction when there is a need not only to obtain a subsample of the CFs table but also to ensure a minimum frequency of taxa in the selection. Such constraints can improve identifiability issues that arise from too few taxa in the hybridization cycle (Solís-Lemus and Ané, 2016; Tiley et al., 2023).

In this study, we showed that it is possible to reduce the dataset without compromising the accuracy of phylogenetic network inference. Reduced datasets allow us to include more taxa in current phylogenetic network analyses and potentially unlock more complex hypotheses on evolutionary histories, including speciation patterns (Mallet, 2007; Fontaine et al., 2015), comparative methods (Bastide et al., 2018), and biogeography (Burbrink and Gehara, 2018). This reduction in data size helps avoid computationally intensive runs, aligning with the principles of green computing (Kumar, 2022). Many avenues remain for applying sparse learning in phylogenetics (e.g., selection of topology-proposal moves), and our study contributes to these efforts. We provide the Qsin algorithm introduced here as a freely available set of scripts with a user-friendly tutorial on GitHub at https://github.com/ulises-rosas/qsin.

## Acknowledgements

The computing for this project was performed at the OU Supercomputing Center for Education & Research (OSCER) at the University of Oklahoma. We thank Mehrnaz Afkhami (Cornell University) for assistance with the creation of Figure 6 and for insightful comments during the development and writing of this project. We are also grateful to Abigail Moore (University of Oklahoma) for helpful comments on the manuscript.

## Code availability

The source code for Qsin is available here: https://github.com/ulises-rosas/qsin.

## Data availability

Both simulated and empirical datasets used throughout this study can be found here: https://github.com/ulises-rosas/qsin

