## Supplementary Material for "Sparse learning for scalable phylogenetic network inference"

**S.II Qsin time complexity** **5**

**S.IIIQsin speed-up analysis** **12**

**S.IVSupplementary Figures** **12**

**S.V Constrained Elastic Net for accounting taxa frequencies** **21**

Table S.1: Summary of notation used throughout the proofs

| Symbol | Description |
|---|---|
| $T$ | Number of taxa (leaf nodes) in the network |
| $n$ | Number of random phylogenetic networks |
| $M$ | Number of regression trees in the ISLE ensemble |
| $\rho$ | fraction of selected CFs rows used in phylogenetic network reconstruction |
| $H$ | Number of hybridization events |
| $h_i$ | $i^{th}$ hybridization event |
| $\mathbf{X}$ | Predictor matrix used in regression |
| $\mathbf{y}$ | Response vector used in regression |
| $\Lambda$ | Set containing $k$ sparsity parameter values |
| $\lambda_k$ | $k^{th}$ sparsity parameter value |
| $\tilde{\lambda}$ | birth rate in the birth-death-hybridization process |
| $\tau$ | Hitting time of a particular state |
| $N_t$ | State of a discrete-time process at time $t$ |
| $r_i$ | Net rate of change at state $i$ |

### S.I SNaQ time complexity

Before analyzing the time complexity of SNaQ, we can state two useful claims that provide a firmer foundation for the analysis:

**Claim 1.** *If a network is level-1, then at most one hybrid edge can be added to any given tree edge of a displayed tree from the network.*

*Proof.* Assume, for contradiction, that a level-1 network allows at least two hybrid edges to be added to the same tree edge of a displayed tree. By minimal counterexample, suppose exactly two such hybrid edges exist. This necessarily leads to a contradiction with the definition of a level-1 network.

Let the tree edge be $e_t$, corresponding to the node pair $(t-1, t)$. Let $h_i$ denote the $i^{\text{th}}$ hybrid node, and let $p_{h_i}$ denote the one parent of $h_i$. We consider the four possible scenarios in which two hybrid edges could lie on the same tree edge:

**Case 1** ($h_1$ and $h_2$ added)**.** *Without loss of generality, assume that the hybrid nodes $h_1$ and $h_2$ are inserted along $e_t$ from top to bottom, subdividing $e_t$ into three segments:*

$$\begin{aligned} e_{h_1} &= (t-1, h_1), \\ e_{h_2} &= (h_1, h_2), \\ e_{t_u} &= (h_2, t). \end{aligned}$$

*Here, $e_{h_1}$ appears in both the $h_1$-based and $h_2$-based hybridization cycles. This contradicts the level-1 assumption, which states that each edge can be part of at most one cycle. We follow the similar rationale for the rest of the cases.*

**Case 2** ($p_{h_1}$ and $p_{h_2}$ added)**.** *Without loss of generality, assume that $p_{h_1}$ and $p_{h_2}$ are inserted along $e_t$ from top to bottom, subdividing it into:*

$$e_{p_{h_1}} = (t-1, p_{h_1}), \quad e_{p_{h_2}} = (p_{h_1}, p_{h_2}), \quad e_{t_u} = (p_{h_2}, t).$$

*Again, $e_{p_{h_1}}$ appears in both the $h_1$-based and $h_2$-based hybridization cycles, violating the level-1 assumption.*

**Case 3** ($h_1$ and $p_{h_2}$ added)**.** *Without loss of generality, assume that $h_1$ and $p_{h_2}$ are inserted along $e_t$ from top to bottom, subdividing it into:*

$$e_{h_1} = (t-1, h_1), \quad e_{p_{h_2}} = (h_1, p_{h_2}), \quad e_{t_u} = (p_{h_2}, t).$$

*Here, $e_{h_1}$ is shared between the cycles associated with $h_1$ and $h_2$, again contradicting the level-1 assumption.*

**Case 4** ($p_{h_1}$ and $h_2$ added)**.** *Without loss of generality, assume that $p_{h_1}$ and $h_2$ are inserted along $e_t$ from top to bottom, resulting in:*

$$e_{p_{h_1}} = (t-1, p_{h_1}), \quad e_{h_2} = (p_{h_1}, h_2), \quad e_{t_u} = (h_2, t).$$

*In this case, $e_{p_{h_1}}$ appears in both the $h_1$- and $h_2$-based cycles, which again violates the level-1 constraint.*

In all four cases, we arrive at a contradiction of the level-1 network assumption. Therefore, no more than one hybrid edge can be added to any given tree edge in a level-1 network. □

**Claim 2.** *Let $T$ be the number of taxa and $H$ the number of hybridization events. If a network is level-1 in SNaQ, then $T > H$ in any displayed tree from the network.*

*Proof.* Assume, for contradiction, that the network is level-1 in SNaQ and that $T \leq H$. Then we can derive the following inequalities:

$$H \geq T > T - \frac{3}{2} \geq \left\lfloor \frac{2T-3}{2} \right\rfloor.$$

Thus, we must have

$$H > \left\lfloor \frac{2T-3}{2} \right\rfloor,$$

which implies

$$H = \left\lfloor \frac{2T-3}{2} \right\rfloor + m$$

for some integer $m \geq 1$.

In each hybridization event, a new hybrid node is added, which must have two distinct parents inserted into two different edges. If both parents were added to the same edge, then the entire hybridization cycle would lie within a single tree edge. However, such cases are ignored by SNaQ as they are unidentifiable (Solís-Lemus and Ané, 2016). Therefore, to support $H$ hybridization events, at least $2H$ distinct edges are needed.

But the displayed tree contains only $2T-3$ edges. So to accommodate $H = \lfloor \frac{2T-3}{2} \rfloor + m$ hybridization events, we would need

$$2H = (2T-3) + 2m$$

edges, which exceeds the number available in the tree. By the pigeonhole principle, this implies that at least one edge must contain two or more hybridization events.

To illustrate this, for example, let $m = 1$, and consider the following setup, where the bottom row lists all the edges in the displayed tree, and the rows above indicate the hybrid events they support:

| $h_{\lfloor \frac{2T-3}{2} \rfloor + 1}$ $h_1$ | $h_2$ | $\dots$ | $h_{\lfloor \frac{2T-3}{2} \rfloor}$ | $h_1$ | $h_2$ | $\dots$ | $h_{\lfloor \frac{2T-3}{2} \rfloor}$ | $h_{\lfloor \frac{2T-3}{2} \rfloor + 1}$ |
|---|---|---|---|---|---|---|---|---|
| $e_1$ | $e_2$ | $\dots$ | $e_{\lfloor \frac{2T-3}{2} \rfloor}$ | $e_{\lfloor \frac{2T-3}{2} \rfloor + 1}$ | $e_{\lfloor \frac{2T-3}{2} \rfloor + 2}$ | $\dots$ | $e_{(2T-3)-1}$ | $e_{(2T-3)}$ |

Here, edge $e_1$ supports both $h_1$ and $h_{\lfloor \frac{2T-3}{2} \rfloor + 1}$ hybridization events, which means it contains two hybrid edges—implying two overlapping hybridization cycles.

By the contrapositive of Claim 1: if at least two hybrid edges lie on the same tree edge in a displayed tree, then the network is not level-1. Therefore, this configuration contradicts our original assumption that the network is level-1. □

#### S.I.1 Objective function in SNaQ

Algorithm S.I describes the objective function evaluation, which corresponds to the log-pseudolikelihood score defined in Eq. 1 in the main text. Thus, each evaluation of the objective function requires $O(T^5)$ operations in total, which comes from repeating $O(T^4)$ times step 2a in Algorithm S.I.

**Algorithm S.I** SNaQ objective function evaluation

1. **Input:** a fixed network topology and a vector of parameters $\mathbf{x}$, which includes internal branch lengths and inheritance probabilities.
2. For $j = 1$ to $\binom{T}{4}$:
   (a) Extract the subnetwork induced by the quartet $j$: $O(T)$ operations
   (b) Using $\mathbf{x}$, compute the quartet $j$ log-likelihood score: $O(1)$ operations
3. Aggregate all quartet log-likelihoods into an overall log-pseudolikelihood score: $O(T^4)$ operations

#### S.I.2 Parameter optimization in SNaQ

Now, to obtain the log-pseudolikelihood score for a given network topology, it is necessary to optimize both the internal branch lengths and the inheritance probabilities. SNaQ accomplishes this using a derivative-free optimization algorithm called BOBYQA (see Algorithm S.II) (Powell et al., 2009).

Let $m$ denote the total number of parameters to be optimized—including both branch lengths and inheritance probabilities—and let $d$ denote the number of iterations of the algorithm. A quadratic approximation of the objective function is constructed by evaluating the objective function (see Section S.I.1) approximately $m$ times to numerically approximate its gradient.

**Algorithm S.II** SNaQ parameter optimization using BOBYQA

1. Initialize the parameter vector $\mathbf{x} \in \mathbb{R}^m$; let $F(\mathbf{x})$ be the SNaQ objective function (see Algorithm S.I); let $\mathbf{e}_j \in \mathbb{R}^m$ denote a standard basis vector (i.e., a vector of zeros with a 1 in the $j$-th component); and let $h$ be a constant defined by the BOBYQA algorithm.
2. For $i = 1$ to $m$:
   (a) Define perturbations $\mathbf{x} + h\mathbf{e}_i$ and $\mathbf{x} - h\mathbf{e}_i$: $O(m)$ operations
   (b) Evaluate $F(\mathbf{x} + h\mathbf{e}_i)$ and $F(\mathbf{x} - h\mathbf{e}_i)$: $O(2T^5)$ operations
   (c) Approximate the gradient component:

$$g_i = \frac{F(\mathbf{x} + h\mathbf{e}_i) - F(\mathbf{x} - h\mathbf{e}_i)}{2h}$$

   $O(1)$ operations
3. Construct the initial quadratic model $Q_1$ using the approximated gradients: $O(m^2)$ operations
4. For $j = 1$ to $d$:
   (a) Solve the quadratic problem $Q_j$ to update $\mathbf{x}$: $O(m^2)$ operations
   (b) Evaluate the objective function at the new interpolation point: $O(T^5)$ operations
   (c) Update the quadratic model $Q_j$: $O(m^2)$ operations

The most computationally expensive components of parameter optimization are outlined in Algorithm S.II and considering the most dominant components for each step in it, we find:

$$\underbrace{O(m)}_{\text{Step 1}} + \underbrace{O\left(mT^5\right)}_{\text{Step 2}} + \underbrace{O\left(m^2\right)}_{\text{Step 3}} + \underbrace{O(d) \times \left[O\left(2m^2\right) + O\left(T^5\right)\right]}_{\text{Step 4}} \leq O\left(2mT^5\right) + O\left(3dm^2\right) + O\left(dT^5\right)$$
$$= O\left(mT^5\right) + O\left(m^2\right) + O\left(T^5\right)$$

If we are creating networks with $T$ taxa, then the number of internal branches in the unrooted displayed tree is $T-3$. Let $H$ denote the number of hybridization events. Since each hybridization event involves approximately three edges, then we can assume $m \approx T + 3H$. By Claim 2 which states that $T > H$, the above simplifies as:

$$\begin{aligned} O\left(mT^5\right) + O\left(m^2\right) + O\left(T^5\right) &= O\left(\{T+3H\}T^5\right) + O\left(\{T+3H\}^2\right) + O\left(T^5\right) \\ &\leq O\left(4T \times T^5\right) + O\left(16T^2\right) + O\left(T^5\right) \\ &= O\left(4T^6\right) + O\left(16T^2\right) + O\left(T^5\right) \\ &\leq O\left(21T^6\right) \\ &= O\left(T^6\right) \end{aligned} \tag{S.1}$$

Therefore, the overall time complexity of optimizing the branch lengths and inheritance probabilities with SNaQ using BOBYQA is $O(T^6)$.

#### S.I.3 Heuristic search

Finally, the most computationally dominant steps of the SNaQ heuristics are found in Algorithm S.III. Let $D$ be the total number of iterations; then the total time complexity of SNaQ is $O(DT^6)$. If the algorithm visits at least all the edges as part of the hill-climbing search, then $D \geq T$. Assuming $D = T$, the total time complexity of SNaQ becomes $O(T^7)$.

**Algorithm S.III** SNaQ heuristics

Repeat until the stopping criterion is met:

1. Copy the current network: $O(T)$ operations.
2. Perform a move and update attributes: $O(1)$ operations.
3. Optimize network parameters (e.g., branch lengths; see Algorithm S.II): $O(T^6)$ operations.

We arbitrarily assumed that the number of moves per run is in the order of T so that we can upper bound SNaQ's time complexity. This is similar to how Neighbor Joining gradually constructs the phylogenetic tree edge by edge (Yang, 2014). However, we also conducted simulations to evaluate how well this assumption holds in practice (Fig. S.1), and we found that it seems reasonable. Although it is a theoretical assumption, empirically, it is not too far off.

### S.II Qsin time complexity

Before analyzing the time complexity of the Qsin pipeline presented in Algorithm 3—which is developed through six propositions and one main theorem—we first state a useful claim that provides a foundation for the analysis:

**Claim 3.** *Let $\tilde{\lambda}$, $\mu$, $\nu_+$, $\nu_-$ be the birth and death rates, and the generative and degenerative hybridization rates, respectively. If $\tilde{\lambda} > \mu$ and $\nu_+ > \nu_-$, then, on average, the SiPhyNetwork*

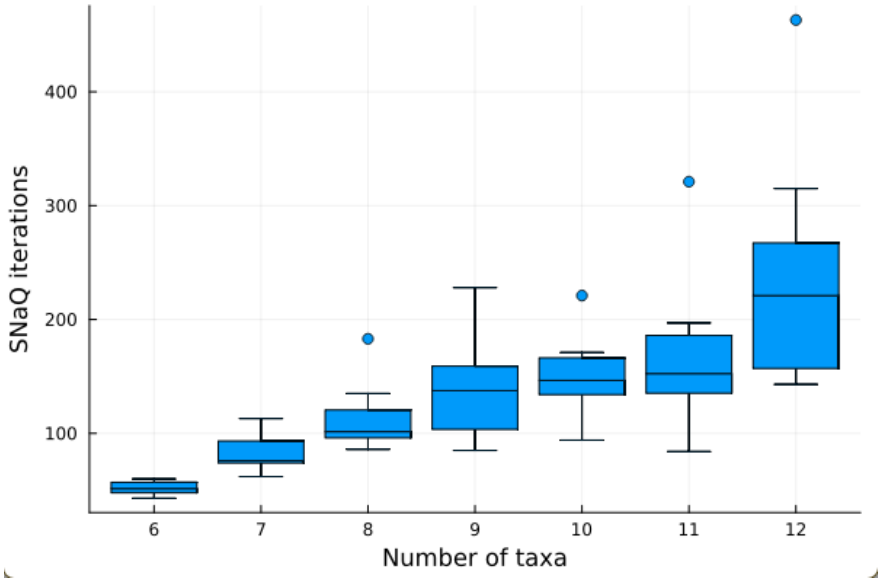

Figure S.1: The number of iterations per run approximately increases linearly as the number of species increases in simulated datasets. Simulations conducted using version 1.0 of the SNaQ Julia package with default arguments. True networks were generated at random by (1) simulating a random tree with the given number of taxa from a star tree using the PhyloCoalSimulations Julia package (Fogg et al., 2023), before (2) randomly selecting origin and destination edges from which a single reticulation is added into the network, and then finally (3) scaling branch lengths such that the average branch length of the network is 1.0. The concordance factor data used as input to SNaQ was then computed from 250 gene trees that were simulated from each network using PhyloCoalSimulations.

*algorithm, under the birth–death–hybridization process, requires $T$ iterations to generate a network with $T$ taxa.*

*Proof.* Although the time complexity of the algorithm for generating a single network using SiPhyNetworks (Justison et al., 2023) was not explicitly described when the method was proposed, we can estimate it based on established analyses of birth-death process simulation (Huelsenbeck, 2018).

The birth–death–hybridization process is an irreducible Markov process (i.e., all states communicate) whose expected change in the number of lineages per unit time is positive and grows approximately quadratically with the state index $i$. More precisely, the net rate of increase at state $i$ is (Justison and Heath, 2023):

$$r_i = \tilde{\lambda}_i - \mu_i = i(\tilde{\lambda} - \mu) + \binom{i}{2}(\nu_+ - \nu_-).$$

Since by assumption $\tilde{\lambda} > \mu$ and $\nu_+ > \nu_-$, then $r_i > 0$ for all $i$. When $r_i > 0$ for all $i$, the process is transient (Lefebvre, 2007, pp. 141) and exhibits a systematic drift toward larger states. Let $\nu_{i,i}$ denote the total jump rate out of state $i$, defined as:

$$\nu_{i,i} = \tilde{\lambda}_i + \mu_i = i(\tilde{\lambda} + \mu) + \binom{i}{2}(\nu_+ + \nu_-),$$

Let $N_t$ be the random variable describing the state of the algorithm at time step $t \in \{0, 1, 2, \dots\}$. Then, the one-step mean drift of the discrete-time chain is given by:

$$\mathbb{E}[N_{t+1} - N_t \mid N_t = i] = (-1)p_{i,i-1} + (0)p_{i,i} + (1)p_{i,i+1} = \frac{r_i}{\nu_{i,i}} = \frac{2(\tilde{\lambda} - \mu) + (i-1)(\nu_+ - \nu_-)}{2(\tilde{\lambda} + \mu) + (i-1)(\nu_+ + \nu_-)}.$$

This expression converges to $\frac{\nu_+ - \nu_-}{\nu_+ + \nu_-}$ as $i \to \infty$. Moreover, by induction (Abbott, 2015, pp. 56), the sequence defined by this expression over all $i$ is monotone. When $i = 1$ (i.e., at $t = 0$), the drift simplifies to $\frac{\tilde{\lambda} - \mu}{\tilde{\lambda} + \mu}$, which is strictly positive. Thus, we have two bounds but do not know which is the smaller. Therefore, for all $i \geq 1$, we can set the lower bound of the mean drift by the minimum of the two:

$$\mathbb{E}[N_{t+1} - N_t \mid N_t = i] \geq \varepsilon := \min\left\{\frac{\tilde{\lambda} - \mu}{\tilde{\lambda} + \mu}, \frac{\nu_+ - \nu_-}{\nu_+ + \nu_-}\right\} > 0.$$

Now, define another random variable $X_t := T - N_t$, where $T$ is the target state at which the algorithm stops. Then the drift inequality becomes:

$$\mathbb{E}[X_t - X_{t+1} \mid X_t = T - i] \geq \varepsilon.$$

Let $\tau$ denote the first hitting time of state $X_t = 0$, which is equivalent to the first time the process reaches state $T$. Then, by the Additive Drift Theorem (He and Yao, 2004; Kötzing, 2014; Lengler, 2019), the expected value of $\tau$ satisfies:

$$\mathbb{E}[\tau] \leq \frac{\mathbb{E}[X_0]}{\varepsilon} = \frac{T-1}{\varepsilon} = O(T).$$

□

**Proposition 1.** *In Algorithm 3, simulating data takes $O(nT^5)$ operations.*

*Proof.* By Claim 3, we know that simulating a network under the birth–death–hybridization process, requires $T$ iterations to generate a network with $T$ taxa.

In each iteration, hybridization, birth, and death events occur with different frequencies. However, all events take constant time since they involve, at most, the creation of three new nodes (in the case of a hybridization event) or simple rearrangements—each of which requires constant time. Therefore, the time complexity per network is $O(1) \times O(T) = O(T)$.

We generate up to $4n$ networks to ensure that at least $n$ of them are level-1 networks, resulting in an overall time complexity of $O(nT)$ for network generation.

For each network, we traverse all the branch lengths to rescale them to a specified average value. A level-1 network contains three branch lengths per hybridization event, giving $3H$, and approximately $2T - 3$ branch lengths from the underlying displayed tree, yielding a total of $O(3H + 2T)$ passes per network. By Claim 2, we know that $T > H$, which implies:

$$O(3H + 2T) \leq O(5T) = O(T)$$

Thus, over $n$ networks, the total complexity for branch length scaling is $O(nT)$.

Finally, evaluating the objective function takes $O(T^5)$ time per network (see Section S.I.1), leading to $O(nT^5)$ time for all networks.

Putting it all together—network generation, branch length transformation, and objective function evaluation—the total time complexity is:

$$O(nT) + O(nT) + O(nT^5) \leq O(3nT^5) = O(nT^5)$$

This dominates the cost of writing and reading the generated matrix $(\mathbf{X}, \mathbf{y})$ (see Eq. 2) to and from memory, which takes $O(nT^4)$ for $n$ observations and $T^4$ columns. □

**Proposition 2.** *In Algorithm 3, fitting the ensemble takes* $O(MT^4\, n \log n)$ *operations.*

*Proof.* The time complexity of constructing a regression tree primarily involves two components (Hastie et al., 2009, pp. 334):

i) Sorting the $p$ feature columns, which takes $O(p\, n \log n)$ time.

ii) Performing the split computations, which takes $O(dnp)$ time for a tree of depth $d$.

In our study, we use shallow trees (small $d$), so the split computation simplifies to $O(np)$. Thus, the dominant cost of constructing the tree becomes the sorting step:

$$O(p\, n \log n) + O(np) \leq O(2p\, n \log n) = O(p\, n \log n)$$

For predictions, evaluating a single observation requires $O(d)$ operations—one per level of the tree. With $n$ observations and shallow trees, the prediction complexity becomes $O(nd) = O(n)$.

These two dominant operations are repeated $M$ times in the ISLE algorithm, so the total time complexity is:

$$O(M) \times [O(p\, n \log n) + O(n)] = O(Mp\, n \log n)$$

Finally, we assume that $p = \frac{1}{3}T^4$, since $p$ corresponds to the number of rows in the concordance factor table, which is on the order of $T^4$. Substituting this into the expression yields the overall complexity:

$$O(M\frac{1}{3}T^4\, n \log n) \leq O(MT^4\, n \log n)$$

□

**Proposition 3.** *In Algorithm 3, post-processing takes* $O(nM)$ *operations.*

*Proof.* We implemented Elastic Net using the naive coordinate descent method (Hastie et al., 2015) and it has a time complexity of $O(np)$, where $n$ is the number of samples and $p$ is the number of predictors. In our setting, the predictors correspond to the $M$ models in the ensemble, i.e., $p = M$. Therefore, the total time complexity for this step is $O(nM)$. □

**Proposition 4.** *In Algorithm 3, row selection takes* $O(nM + \rho T^4)$ *operations.*

*Proof.* We consider two scenarios: one where $\lambda_k$ is pre-specified, and another where it is selected by minimizing test error.

**Case 1** (Selection with a pre-specified $\lambda_k$)**.** *Algorithm 2 contains three main steps inside its loop:*

*i) Checking if a regression tree is inactive, which takes constant time $O(1)$.*

*ii) Extracting the index set from split variables used in the regression tree, which takes $O(d)$ time, where $d$ is the (fixed) depth of the tree.*

*iii) Merging these indices into a global index set. Using a hash table, this takes $O(d)$ in expectation.*

*Each iteration thus takes $O(d)$ time, and the loop runs for $M$ models. Since $d$ is constant, the total time complexity is:*

$$O(M) \times [O(1) + O(d) + O(d)] \leq O(3dM) = O(M)$$

**Case 2** (Selection of $\lambda_k$ via test error minimization)**.** *In Algorithm S.IV, we first generate the matrix $\mathbf{T}_{test} \in \mathbb{R}^{n \times M}$ from unseen test data. As discussed previously, predicting outcomes from $M$ models over $n$ samples requires $O(nM)$ operations.*

*Within the loop over sparsity levels $\lambda_i \in \Lambda$, we compute the weighted prediction $\mathbf{T}_{test}\mathbf{w}^{\lambda_i}$, which also takes $O(nM)$ time. Since the number of sparsity values $|\Lambda| = k$ is constant, the total complexity of this loop is $O(knM) = O(nM)$. Other operations (e.g., error comparison and assignment) are negligible in comparison.*

*According to Case 1, generating the index set $I^{\lambda_k}$ takes $O(M)$. Therefore, the overall complexity for this case is:*

$$O(nM) + O(M) \leq O(2nM) = O(nM)$$

**Algorithm S.IV** Select $\lambda_k$ by minimizing test error

```
Generate matrix T_test from unseen (test) data as described in Section 2.4.
Initialize: λ_k ← ∅, ε_k ← ∞
for λ_i ∈ Λ do
    Compute prediction error on test data:
        ε_i = sqrt( (1/n) ||y_test − T_test w^{λ_i}||_2^2 )
    if ε_i < ε_k then
        λ_k ← λ_i
        ε_k ← ε_i
    end if
end for
return λ_k
```

$$\varepsilon_i = \sqrt{\frac{1}{n}\left\|\mathbf{y}_{\text{test}} - \mathbf{T}_{\text{test}}\mathbf{w}^{\lambda_i}\right\|_2^2}$$

The worst-case scenario in terms of computational cost occurs when $\lambda_k$ is selected via error minimization. Thus, we take $O(nM)$ as the complexity of this part.

Finally, writing the selected indices $I^{\lambda_k}$ to a file involves looping through all selected features. Since these correspond to a fraction $\rho \in (0, 1]$ of the $T^4$ total rows in the concordance factor table, this step adds $O(\rho T^4)$ operations. Hence, the overall complexity for row selection is: $O(nM + \rho T^4)$ □

**Proposition 5.** *Cross-validation for the ISLE algorithm takes $O(MT^4 n \log n)$ operations.*

*Proof.* To perform cross-validation, it is necessary to first streamline the fitting process—which includes both ensemble generation and post-processing via Elastic Net—in order to evaluate a given hyperparameter set on $l$-folds of the data. Let $n_l$ denote the approximate number of observations in each fold. Then, by Propositions 2 and 3, the total number of operations required to fit a hyperparameter set on $n_l$ observations is

$$O(MT^4 n_l \log n_l + Mn_l).$$

By Proposition 4, Case 2, adding the cost of error evaluation, this becomes

$$O(MT^4 n_l \log n_l + 2Mn_l).$$

Let $c$ be the total number of hyperparameter sets to evaluate on each of the $l$-folds. This results in a total of $cl$ evaluations. Note that $cl$ is a positive constant since $l = 5$ and $c$ is capped at 100 due to the use of a random sampling strategy for parameter combinations (Bergstra and Bengio, 2012). Therefore, the total computational complexity for fitting and evaluation error during cross-validation is:

$$\begin{aligned} O\left(cl \cdot \{MT^4 n_l \log n_l + 2Mn_l\}\right) &\leq O\left(cl \cdot \{MT^4 n \log n + 2Mn\}\right) \\ &\leq O(cl \cdot 3MT^4 n \log n) \\ &= O(MT^4 n \log n). \end{aligned}$$

We store all error vectors—where each item corresponds to the error for a given sparsity parameter $\lambda_k$—in a linked list, so the total cost of appending them into the list is $O(cl)$.

Next, we use a hash table with keys corresponding to hyperparameter sets to compute the average error across all folds. For each key, this requires sum vectors of length $k$ (the number of $\lambda_k$ values in the Elastic Net path), and this operation is repeated $cl$ times. Thus, the total complexity for computing the fold-average errors is: $O(clk)$.

We then make one pass over the hash table to identify the key (i.e., the hyperparameter set) with the lowest average error. This is similar to what is done in Algorithm S.IV. For each iteration, finding the minimum in the error vector takes $O(k)$, and since there are $l$ folds, the total complexity of this process is $O(lk)$.

Putting everything together—including fitting and error evaluation—the total complexity of cross-validation for selecting the best hyperparameter set is:

$$\begin{aligned} O(MT^4 n \log n) + O(cl) + O(clk) + (lk) &\leq O(MT^4 n \log n) + O(3clk) \\ &= O(MT^4 n \log n) + O(1) \\ &\leq O(2MT^4 n \log n) \\ &= O(MT^4 n \log n). \end{aligned}$$

□

**Proposition 6.** *If we assume the number of SNaQ iterations is at most on the order of $T$, then, in Algorithm 3, reconstructing the network takes $O(\rho T^7)$ operations.*

*Proof.* We essentially adopt the same assumptions outlined in Section S.I.2. Specifically, the parameter optimization from Eq. S.1 can be rewritten as:

$$\begin{aligned} O\left(m\rho T^5\right) + O\left(m^2\right) + O\left(\rho T^5\right) &= O\left(\{T+3H\}\rho T^5\right) + O\left(\{T+3H\}^2\right) + O\left(\rho T^5\right) \\ &\leq O\left(4T \times \rho T^5\right) + O\left(16T^2\right) + O\left(\rho T^5\right) \\ &= O\left(4\rho T^6\right) + O\left(16T^2\right) + O\left(\rho T^5\right) \\ &\leq O\left(21\rho T^6\right) \\ &= O\left(\rho T^6\right) \end{aligned} \tag{S.2}$$

Assuming the number of optimization iterations is at most $T$, the total number of operations for this step is:

$$\underbrace{O(\rho T^6)}_{\substack{\text{Parameter} \\ \text{optimization}}} \times \underbrace{O(T)}_{\substack{\text{Number of} \\ \text{iterations}}} = O(\rho T^7)$$

□

**Theorem 1** (Overall Time Complexity of Algorithm 3)**.** *Let $T$ be the number of taxa, $n$ be the number of simulations, $M$ the number of used regression trees, and $\rho \in (0,1]$ the fraction of selected CFs rows used in reconstruction. Assume the number of SNaQ iterations is at most on the order of $T$. Then, the total time complexity of Algorithm 3 is $O\left(nT^5 + MT^4 n \log n + \rho T^7\right)$.*

*Proof.* We count the operations contributed by each step:

$$\begin{aligned} O\left(nT^5\right) &\quad \text{(Simulate data, Proposition 1)} \\ + O\left(MT^4\, n \log n\right) &\quad \text{(Cross-validation, Proposition 5)} \\ + O\left(MT^4\, n \log n\right) &\quad \text{(Fit ensemble, Proposition 2)} \\ + O(nM) &\quad \text{(Post-process, Proposition 3)} \\ + O(nM + \rho T^4) &\quad \text{(Row selection, Proposition 4)} \\ + O(\rho T^7) &\quad \text{(Reconstruct network, Proposition 6)} \end{aligned}$$

Combining and simplifying these terms:

$$\begin{aligned} O(nT^5) + O(2MT^4 n \log n + 2nM) + O(\rho T^4 + \rho T^7) &\leq O(nT^5) + O(4MT^4 n \log n) + O(2\rho T^7) \\ &= O(nT^5) + O(MT^4 n \log n) + O(\rho T^7) \\ &= O\left(nT^5 + MT^4 n \log n + \rho T^7\right). \end{aligned}$$

□

### S.III Qsin speed-up analysis

**Corollary 1.** *Let $T$ be the number of taxa. Under the same assumptions outlined in Theorem 1, and as $T \to \infty$, the worst-case scenario yields a speed-up factor of $O(1)$ (i.e., no speed-up) and the best-case scenario yields a speed-up factor of $O(T^3)$ relative to SNaQ.*

*Proof.* Under the same assumptions outlined in Theorem 1, the asymptotic complexity of the SNaQ algorithm is $O(T^7)$ (see Supplementary Material S.I for details). We now consider two extreme cases.

**Case 1** (Worst-case scenario)**.** *Suppose the entire dataset is selected, which is on the order of $T^4$. Then,*

$$\rho = O\left(\frac{T^4}{T^4}\right) = O(1).$$

*In this case, the runtime complexity from Theorem 1 becomes*

$$O(nT^5 + T^4 Mn \log n + \rho T^7) = O(nT^5 + T^4 Mn \log n + T^7).$$

*The corresponding speed-up factor as a function of $T$ is*

$$\lim_{T\to\infty} \frac{T^7}{nT^5 + T^4 Mn \log n + T^7} \leq \lim_{T\to\infty} \frac{nT^5 + T^4 Mn \log n + T^7}{nT^5 + T^4 Mn \log n + T^7} = O(1).$$

*Hence, there is essentially no speed-up.*

**Case 2** (Best-case scenario)**.** *Suppose the algorithm selects only the minimum number of rows required to cover all species. Since each row contains four species, this number is approximately $\lceil T/4 \rceil$. In this case,*

$$\rho = O\left(\frac{\lceil T/4 \rceil}{T^4}\right) = O\left(\frac{1}{T^3}\right).$$

*The runtime complexity then becomes*

$$O(nT^5 + T^4 Mn \log n + \rho T^7) = O(nT^5 + T^4 Mn \log n + T^4).$$

*The resulting speed-up factor is*

$$\lim_{T\to\infty} \frac{T^7}{nT^5 + T^4 Mn \log n + T^4} = \lim_{T\to\infty} \frac{1}{\frac{nT}{T^3} + \frac{Mn \log n}{T^3} + \frac{1}{T^3}} \leq \lim_{T\to\infty} \frac{1}{3/T^3} = O(T^3).$$

Therefore, the overall speed-up lies between $O(1)$ (Case 1) and $O(T^3)$ (Case 2). □

### S.IV Supplementary Figures

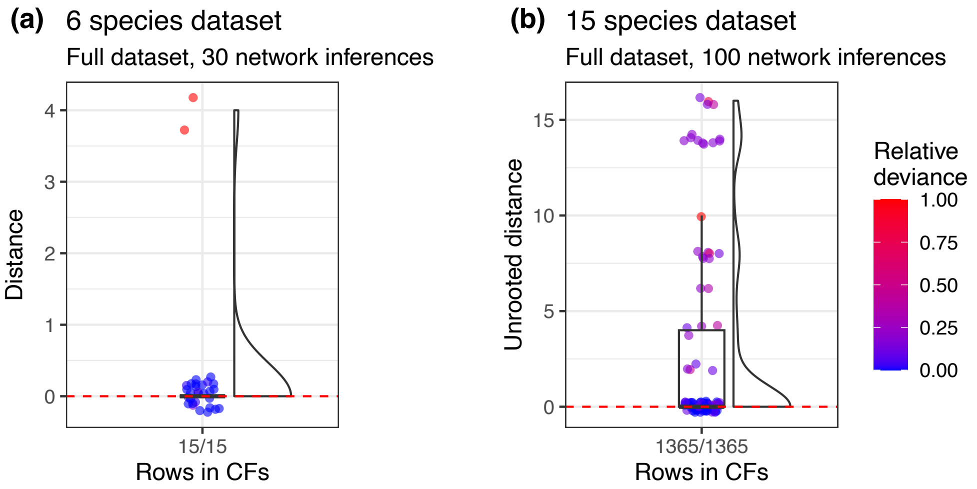

Figure S.2: Accuracy using all rows of the CFs table for both simulated datasets. **(a)** shows 93.3% accuracy for the first simulated dataset, where each repetition used 10 independent runs. **(b)** shows 69% accuracy for the second simulated dataset. Relative deviance denotes the normalized log-pseudolikelihood score.

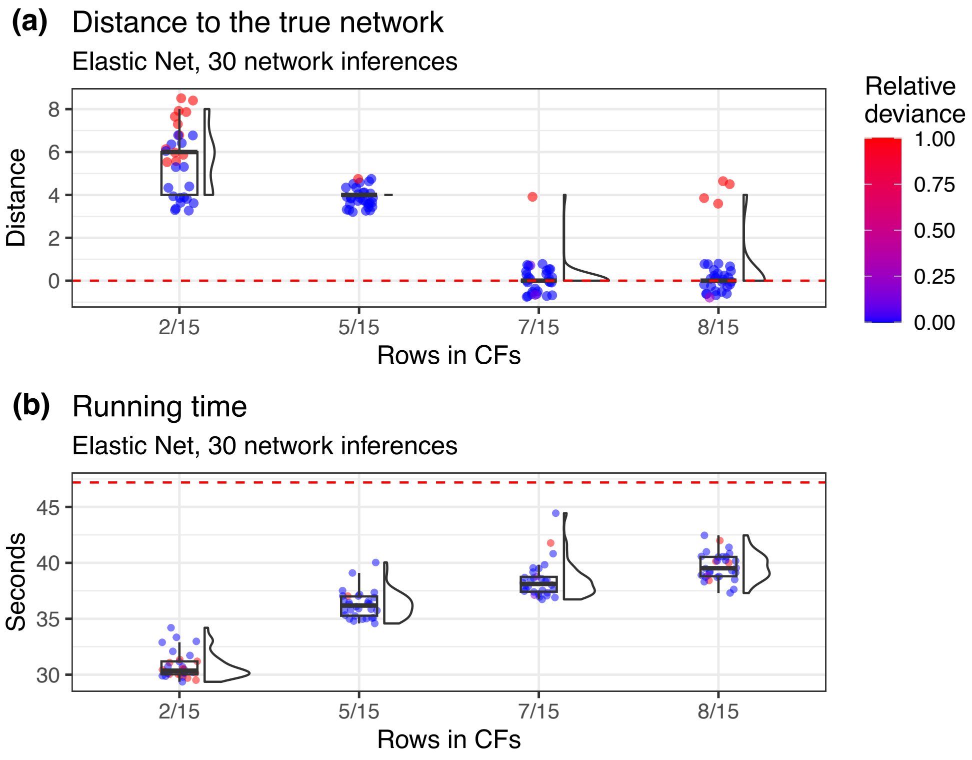

Figure S.3: Accuracy of reconstruction using subsampling guided by Elastic Net when branch lengths come from an exponential distribution draw with an average of 1.25 on a CFs table with 15 rows. **(a)** shows the distance to the true network. With 8 rows, the true network is recovered 86% of the 30 repetitions. The red dashed line indicates zero distance. **(b)** shows inference time for different row selection sizes. The red dashed line indicates the median time required to reconstruct the network using the full dataset, which is approximately 47.2 seconds. Each repetition used 10 independent runs. Relative deviance denotes the normalized log-pseudolikelihood score.

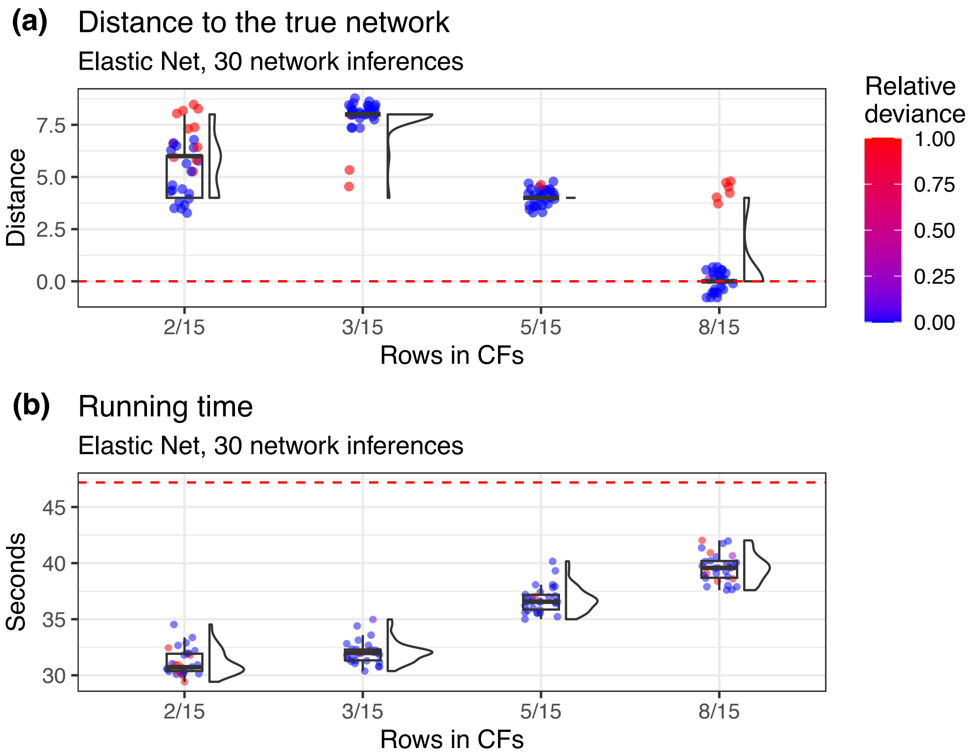

Figure S.4: Accuracy of reconstruction using subsampling guided by Elastic Net when branch lengths are not modified and come directly from the SiPhyNetworks on a CFs table with 15 rows. **(a)** shows the distance to the true network. With 8 rows, the true network is recovered 80% of the 30 repetitions. The red dashed line indicates zero distance. **(b)** shows inference time for different row selection sizes. The red dashed line indicates the median time required to reconstruct the network using the full dataset, which is approximately 47.2 seconds. Each repetition used 10 independent runs. Relative deviance denotes the normalized log-pseudolikelihood score.

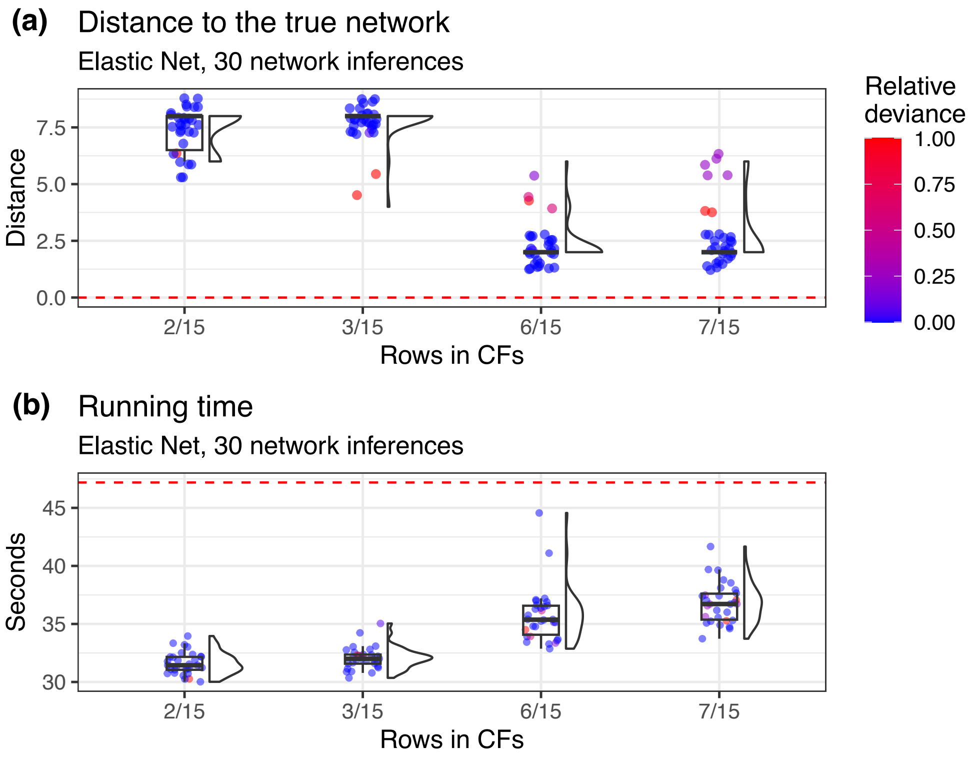

Figure S.5: Accuracy of reconstruction using subsampling guided by Elastic Net when branch lengths of the random network topologies are optimized on a CFs table with 15 rows. **(a)** shows the distance to the true network. With 7 rows, the true network is recovered 0% of the 30 repetitions. The red dashed line indicates zero distance. **(b)** shows inference time for different row selection sizes. The red dashed line indicates the median time required to reconstruct the network using the full dataset, which is approximately 47.2 seconds. Each repetition used 10 independent runs. Relative deviance denotes the normalized log-pseudolikelihood score.

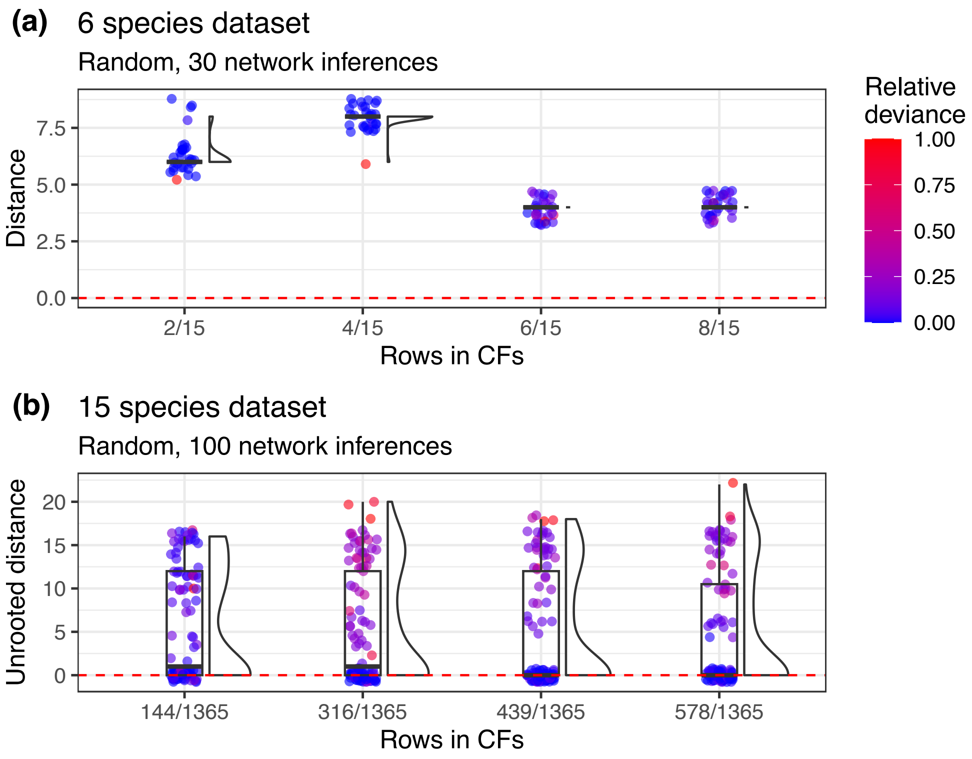

Figure S.6: Accuracy for random selection of rows. **a)** Accuracy of randomly selected rows with the same sizes as in Fig. 2, where each repetition used 10 independent runs. **b)** Accuracy of randomly selected rows with the same size as in Fig. 5. Relative deviance denotes the normalized log-pseudolikelihood score.

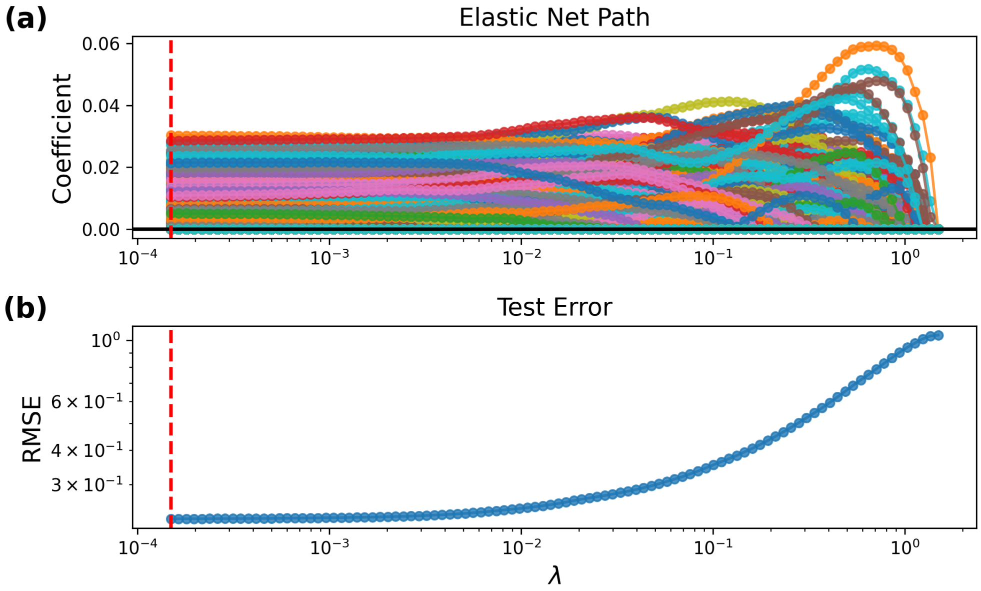

Figure S.7: The Elastic Net path applied to the ISLE ensemble on the *Xiphophorus* dataset, which involves a CFs table with 10,626 rows and 24 species. Hyperparameters were set to $\alpha = 0.5$, 2 leaf nodes, $\eta = 0.1487$, and $\nu = 0.0775$. In **(a)**, each colored line represents a regression tree. From right to left, we observe that when $\lambda$ is at its largest value, no regression trees are selected — as indicated by all colored lines remaining at zero. The red dashed vertical line marks the point along the path with minimun testing error. **(b)** shows the testing RMSE along the elastic net path, reaching its minimun testing error when 788 out of 4,000 regression trees are active—corresponding to 763 rows in the CFs table.

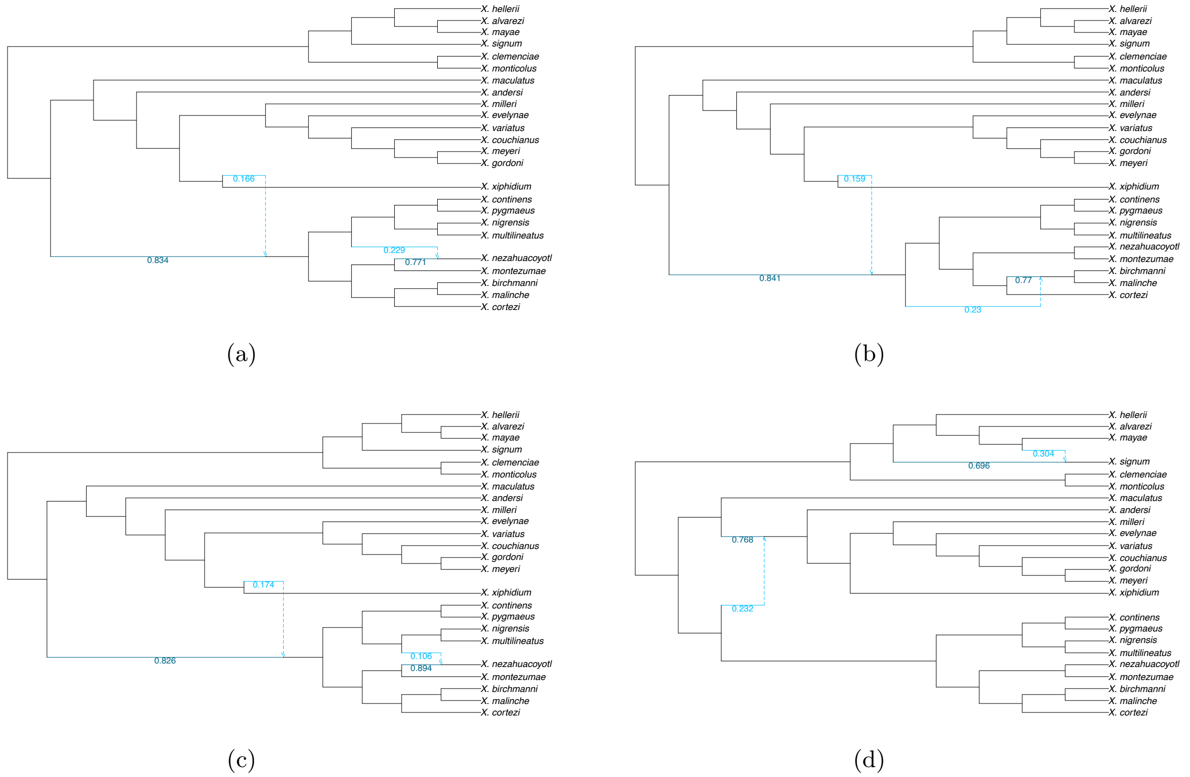

Figure S.8: Phylogenetic network reconstruction for the *Xiphophorus* dataset using the subsample of 763 rows (7.81% of the CFs table), with log-pseudolikelihood scores of **(a)** $-758.770$, **(b)** $-775.370$, **(c)** $-798.085$, and **(d)** $-996.226$. The subsample is identical to that used in Fig. 6.

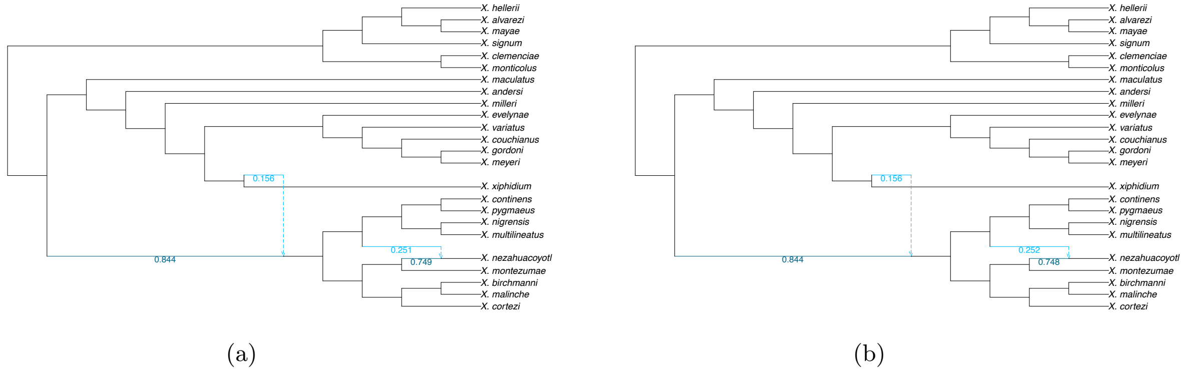

Figure S.9: Phylogenetic network reconstruction for the *Xiphophorus* dataset using the subsample of 763 rows (7.81% of the CFs table), with **(a)** 25 and **(b)** 30 independent runs. The corresponding log-pseudolikelihood scores were $-709.162$ and $-709.158$, respectively. The subsample is identical to that used in Fig. 6.

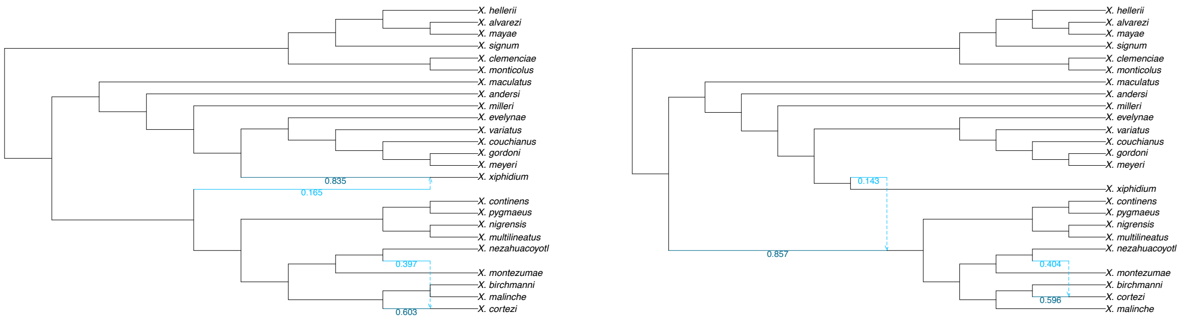

(a) (b)

Figure S.10: Phylogenetic network reconstruction for the *Xiphophorus* dataset using a random subsample of 763 rows (7.81% of the CFs table), with **(a)** 25 and **(b)** 30 independent runs. The corresponding log-pseudolikelihood scores were −908.548 and −795.571, respectively.

### S.V Constrained Elastic Net for accounting taxa frequencies

In Eq. 3, we can constrain $\mathbf{w}$ to take values between 0 and 1 with the following condition:

$$\mathbf{1}^\top \mathbf{w} = 1 \quad \text{and} \quad \mathbf{w} \geq \mathbf{0},$$

where $\mathbf{0} \in \mathbb{R}^M$ and $\mathbf{1} \in \mathbb{R}^M$ are column vectors of zeros and ones, respectively. This is a common technique in stacked generalization, where $\mathbf{w}$ can be interpreted as the posterior model probabilities for the ensemble (Hastie et al., 2009, pp. 290). When $\mathbf{w}$ takes values in $[0, 1]$, the weighted sum $\mathbf{z}_t^\top \mathbf{w}$ can be interpreted as the weighted average frequency of taxon $t$, with the weights given by $\mathbf{w}$.

To enforce a minimum frequency, we add the linear constraint:

$$\mathbf{z}_t^\top \mathbf{w} \geq c \quad \text{for all} \quad t \in \{1, \dots, T\},$$

which ensures that the average frequency of each taxon among the $T$ taxa is at least $c$. For convenience, let $\mathbf{Z} = [\mathbf{z}_1, \dots, \mathbf{z}_T]^\top \in \mathbb{R}^{T \times M}$ denote the matrix whose rows are the vectors $\mathbf{z}_t^\top$ for $t \in \{1, \dots, T\}$. With this notation, we can then incorporate the above constraints into Eq. 3 using the following equality and inequality constraints:

$$\begin{aligned} &\text{minimize} \quad \frac{1}{n}\|\mathbf{y} - \mathbf{T}\mathbf{w}\|_2^2 + \lambda \left\{ \frac{1}{2}(1-\alpha)\|\mathbf{w}\|_2^2 + \alpha \|\mathbf{w}\|_1 \right\} \\ &\text{subject to} \quad \mathbf{1}^\top \mathbf{w} = 1 \quad \text{and} \quad \begin{bmatrix} \mathbf{I} \\ \mathbf{Z} \end{bmatrix} \mathbf{w} \geq \begin{bmatrix} \mathbf{0} \\ \mathbf{c} \end{bmatrix}, \end{aligned}$$

where $\mathbf{c} \in \mathbb{R}^T$ is a column vector containing the desired minimum weighted averages for each taxon, and $\mathbf{I} \in \mathbb{R}^{M \times M}$ is the identity matrix. Multiplying both sides of the inequality constraint by $-1$ yields the standard form of the constrained Lasso, for which the entire solution path can be efficiently computed, including the Elastic Net case (see Eq. 6 in Gaines et al., 2018).
